# Heterogeneous progenitor cell behaviors underlie the assembly of neocortical cytoarchitecture

**DOI:** 10.1101/494088

**Authors:** Alfredo Llorca, Gabriele Ciceri, Robert Beattie, Fong K. Wong, Giovanni Diana, Eleni Serafeimidou, Marian Fernández-Otero, Carmen Streicher, Sebastian J. Arnold, Martin Meyer, Simon Hippenmeyer, Miguel Maravall, Oscar Marín

## Abstract

The cerebral cortex contains multiple hierarchically organized areas with distinctive cytoarchitectonical patterns, but the cellular mechanisms underlying the emergence of this diversity remain unclear. Here, we have quantitatively investigated the neuronal output of individual progenitor cells in the ventricular zone of the developing mouse neocortex using a combination of methods that together circumvent the biases and limitations of individual approaches. We found that individual cortical progenitor cells show a high degree of stochasticity and generate pyramidal cell lineages that adopt a wide range of laminar configurations. Mathematical modelling these lineage data suggests that a small number of progenitor cell populations, each generating pyramidal cells following different stochastic developmental programs, suffice to generate the heterogenous complement of pyramidal cell lineages that collectively build the complex cytoarchitecture of the neocortex.

## INTRODUCTION

The mammalian cerebral cortex contains a wide diversity of neuronal types heterogeneously distributed across layers and regions. The most abundant class of neurons in the cerebral cortex are excitatory projection neurons, also known as pyramidal cells (PCs). In the neocortex, PCs can be further classified into several subclasses with unique laminar distributions, projection patterns and electrophysiological properties (Greig et al., 2013; Jabaudon, 2017; Lodato and Arlotta, 2015), and currently available data suggest that several dozen distinct transcriptional signatures can be distinguished among them (Tasic et al., 2018). The relative abundance of the different types of PCs largely determines the distinct cytoarchitectonical patterns observed across different regions of the mammalian neocortex (Brodmann and Gary, 2006).

The diversity of excitatory neurons emerges from progenitor cells in the ventricular zone (VZ) of the developing neocortex known as radial glial cells (RGCs) (Malatesta et al., 2000; Miyata et al., 2001; Noctor et al., 2001). RGCs divide symmetrically to expand the progenitor pool during early stages of corticogenesis. Subsequently, they undergo asymmetric cell divisions to generate clones of PCs directly or indirectly via intermediate progenitor cells (IPCs) (Lui et al., 2011; Taverna et al., 2014). The characteristic vertical organization of migrating PCs in the developing neocortex led to the “radial unit hypothesis”, which postulates that PCs in a given radial column are clonally related (Rakic, 1988). However, the precise mechanisms through which RGCs generate diverse cytoarchitectonical patterns throughout the neocortex remain to be elucidated.

The most commonly accepted view of cortical neurogenesis is based on the notion that RGCs are multipotent and generate all types of excitatory neurons following an exquisite inside-out temporal sequence (Leone et al., 2008; Molyneaux et al., 2007; Rakic et al., 1994). Consistently, progenitor cells cultured in vitro reproduce the temporal sequence of cortical neurogenesis (Gaspard et al., 2008; Shen et al., 2006), and genetic fate mapping experiments have shown that cortical progenitors identified by the expression of the transcription factors Fezf2 and Sox9 are multipotent in vivo (Guo et al., 2013; Kaplan et al., 2017). In contrast to this view, other studies have suggested the existence of fate-restricted cortical progenitors, which would only generate PCs for certain layers of the neocortex (Franco et al., 2012; Garcia-Moreno and Molnar, 2015). However, the interpretation of these results remains a matter of controversy (Eckler et al., 2015; Gil-Sanz et al., 2015).

Our current framework for understanding cortical neurogenesis largely relies on studies that consider RGCs as a homogeneous population. Consistent with this view, recent clonal analyses of the developing neocortex led to the conclusion that progenitor cell behavior conforms to a deterministic program through which individual RGCs consistently generate the same neuronal output (Gao et al., 2014). This would suggest that variations in the organization of cortical areas would exclusively rely on mechanisms of lineage refinement at postmitotic stages, such as programmed cell death. Alternatively, the absence of detailed quantitated data of individual pyramidal cell lineages or methodological caveats may have prevented the identification of a certain degree of heterogeneity in the neuronal output of individual RGCs.

In this study, we have used three complementary approaches to circumvent some of the intrinsic technical biases associated to each of the previously used methods to systematically investigate the clonal organization of pyramidal cell lineages in the cerebral cortex. Our results provide a detailed quantitative assessment of the neurogenic fate of individual VZ progenitor cells that reveal a large diversity of PC lineage configurations. These findings support a stochastic model of cortical neurogenesis through which a limited number of progenitor cell identities would generate the entire diversity of cytoarchitectonical patterns observed in the neocortex.

## RESULTS

### Retroviral tracing of pyramidal cell lineages

To study the cellular mechanisms underlying the generation of pyramidal cells (PCs) in the neocortex, we analyzed the output and organization of neuronal lineages generated by individual progenitor cells. To this end, we first used replication-deficient retroviral vectors that integrate indiscriminately in mitotic cells but only identify cell lineages with fluorescent proteins following Cre-dependent recombination (Ciceri et al., 2013). To specifically label PC lineages, we injected a very low titer cocktail of conditional reporter retroviruses (*rv::dio-Gfp* and *rv::dio-mCherry*) into the lateral ventricle of *Nex-Cre* mouse embryos (*Neurod6^Cre/+^*), in which Cre expression is confined to postmitotic pyramidal neurons (Goebbels et al., 2006) (Figure 1A). Using this approach, we achieved sparse labelling of individual pallial lineages and avoided introducing a bias in the labeling of progenitor cells in the ventricular zone (VZ) (Cepko et al., 2000).

**Figure 1.**
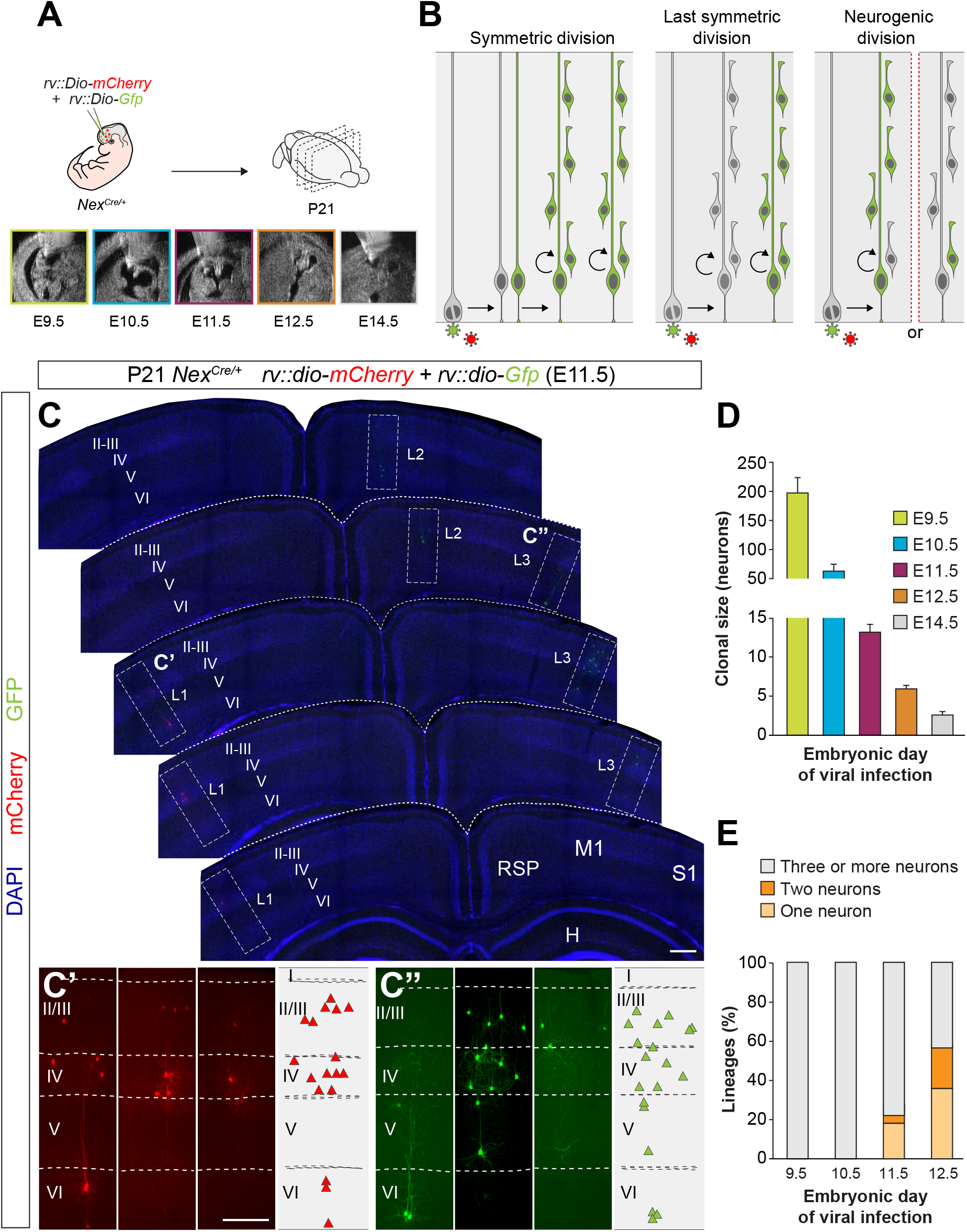
Identification of pyramidal cell lineages with low-titer conditional reporter retroviruses. (A) Experimental paradigm. (B) Schematic representation of the expected labeling outcomes in retroviral lineage tracing experiments. (C) Serial coronal sections through the telencephalon of P21 *Nex^Cre/+^* mice infected with low-titer conditional reporter retroviruses at E11.5. Lineages (L) 1 and 3 are shown at high magnification in C’ and C”, respectively. Dashed lines define external brain boundaries and cortical layers. The schemas collapse lineages spanning across several sections into a single diagram. (D) Quantification of the number of PCs per lineage in P21 *Nex^Cre/+^* mice infected with conditional reporter retroviruses at different embryonic stages. (E) Quantification of the fraction of cortical lineages containing one, two or three or more neurons in P21 *Nex^Cre/+^* mice infected with conditional reporter retroviruses at different embryonic stages. Data are presented as mean ± sem. I–VI, cortical layers I to VI; H, hippocampus area; M1, primary motor cortex; RSD, retrosplenial cortex; S1, primary somatosensory cortex. Scale bars equal 100 μm (C) and 300 μm (C’ and C”).

To identify the developmental stage at which progenitor cells become neurogenic in the cortex, we injected retroviruses at different embryonic days (E9.5 to E14.5) and analyzed the organization of individual PC clusters at postnatal day (P) 21 (Figure 1A). Since a single copy of the viral vector is stably integrated into the host genome, retroviral infection leads to the labelling of only one of the two daughter cells resulting from the division of the infected progenitor cell. Consequently, infection of progenitor cells in the VZ of the pallium labels PC lineages in three main configurations (Figure 1B) depending of the mode of division of the infected progenitor: (i) a large cluster containing more than one lineage, which results from the infection of a self-renewing progenitor cell dividing symmetrically; (ii) a single lineage, which results from the infection of a progenitor cell undergoing its last symmetric division; and (iii) a partial lineage, which results from a neurogenic division of a progenitor cell. In this later case, partial lineages may contain the majority of neurons in the clone, if integration occurs in the progenitor cell, or one or two neurons, if the integration occurs in a neuron or an IPC, respectively.

We observed clusters of neurons with the characteristic morphology of PCs at all stages examined. Systematic mapping at P21 revealed very sparse labelling and widespread distribution of clones throughout the entire neocortex (Figures 1C-C” and Figure S1). The spatial segregation of the lineages was confirmed by the virtual absence of green and red clones within 500 μm of each other in all experiments analyzed (Figure S1). We quantified the number of PCs per clone at P21 following viral infection at different embryonic stages and observed that lineages contain progressively smaller progenies (Figure 1D). This is consistent with the notion that VZ progenitors undergo proliferative symmetric cell divisions early during corticogenesis before they become neurogenic and begin self-renewing via asymmetric divisions (Gotz and Huttner, 2005; Kriegstein and Gotz, 2003). Since neurogenic divisions label one or two neurons in 50% of the cases (Figure 1B), the fraction of one- and two-cell clones found after retroviral infection is indicative of the proportion of neurogenic VZ progenitor cells at each embryonic stage. We observed that these clones represent ~50% of the lineages at E12.5 (Figure 1E). Consistent with previous reports using other methods (Gao et al., 2014), these results indicate that the onset of cortical neurogenesis begins immediately before E12.5, and that at this stage most of VZ progenitor cells are already neurogenic. Thus, we focused subsequent analyses on this stage.

### Heterogeneous neurogenic fate of cortical progenitor cells

We first examined lineages labeled at E12.5 that contained more than two cells, which correspond to the progeny of a VZ progenitor cell (Figure 2A). Consistent with classical models of cortical neurogenesis, we found that most VZ progenitor cells (63%) infected with retroviruses at E12.5 produce translaminar lineages containing neurons in both deep (V and VI) and superficial (II-III and IV) layers of the neocortex (Figures 2B and 2E). However, we also observed that a substantial fraction of lineages in which PCs were confined to either deep (Figures 2C and 2E) or superficial (Figures 2D and 2E) layers (15% and 22%, respectively).

**Figure 2.**
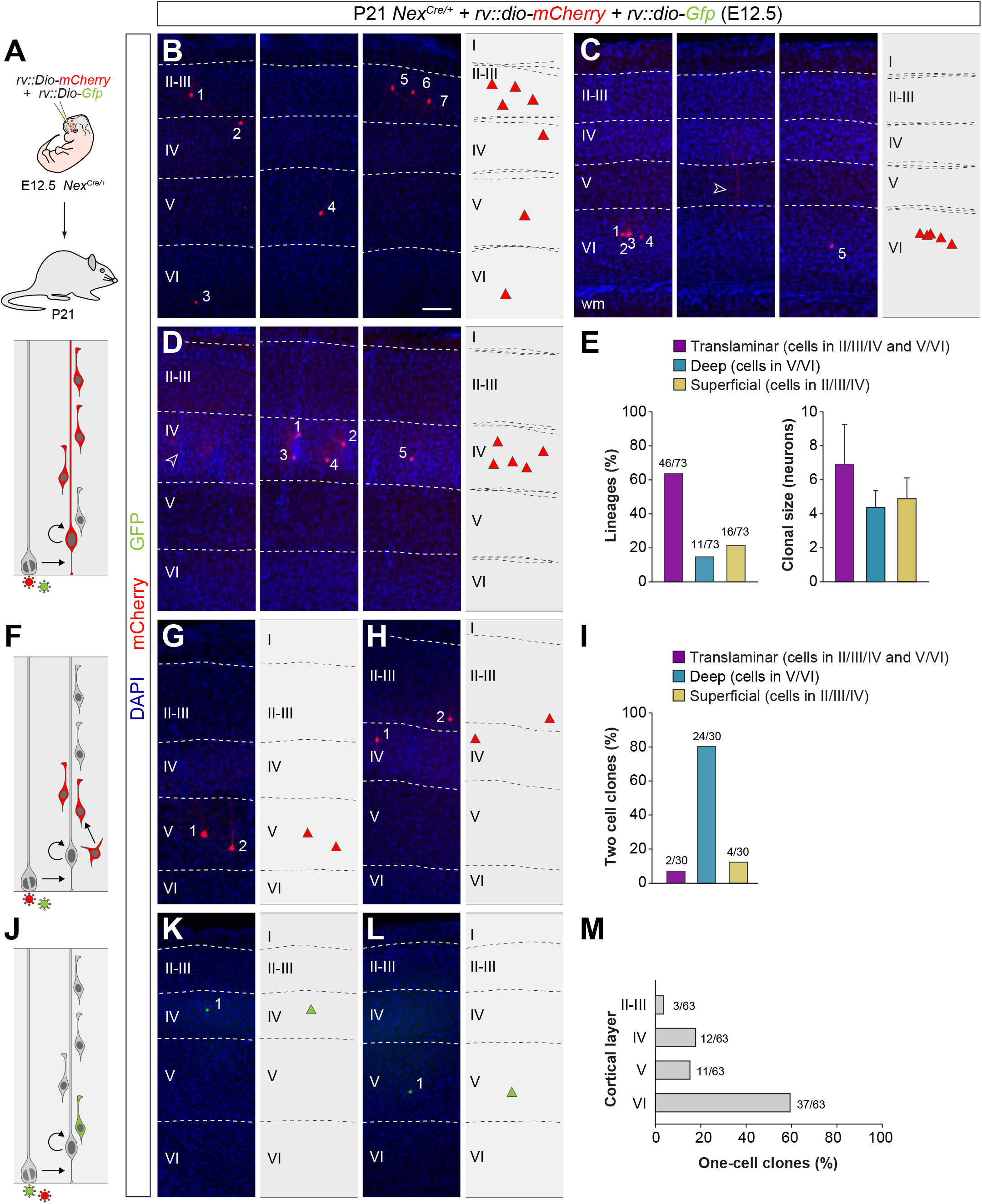
Retroviral-based lineage tracing reveals diverse lineage outcomes. (A) Experimental paradigm. The bottom panel illustrates the expected labeling outcome following retroviral infection of an RGC undergoing a neurogenic cell division in which the viral integration occurs in the self-renewing RGC. (B–D) Serial coronal sections through the cortex of P21 *Nex^Cre/+^* mice infected with low-titer conditional reporter retroviruses at E12.5. The images show examples of translaminar (B), deep-layer restricted (C) and superficial-layer restricted (D) lineages containing three or more cells. Dashed lines define external brain boundaries and cortical layers. The schemas collapse lineages spanning across several sections into a single diagram. (E) Quantification of the fraction of translaminar, deep- and superficial-layer restricted lineages containing three or more cells, and clonal size. Clonal size data are presented as mean ± standard deviation. (F) Expected labeling outcome following retroviral infection of an RGC undergoing a neurogenic cell division in which the viral integration occurs in an IPC (indirect neurogenesis). (G–H) Coronal sections through the cortex of P21 *Nex^Cre/+^* mice infected with low-titer conditional reporter retroviruses at E12.5. The images show examples of superficial and deep layer-restricted two-cell clones. (I) Quantification of the fraction of translaminar, deep and superficial layer-restricted two-cell lineages. (J) Expected labeling outcome following retroviral infection of an RGC undergoing a neurogenic cell division in which the viral integration occurs in a postmitotic neuron (direct neurogenesis). (K–L) Coronal sections through the cortex of P21 *Nex^Cre/+^* mice infected with low-titer conditional reporter retroviruses at E12.5. The images show examples of superficial and deep layer-restricted single-cell clones. (M) Laminar distribution of single-cell clones. Scale bar equals 100 μm.

The distribution of single-cell and two-cell clones following infection of VZ progenitor cells at E12.5 further support the existence of cortical lineages restricted to superficial layers of the neocortex. As expected from the normal progression of neurogenesis in translaminar lineages, most single-cell and two-cell clones (which result from the labeling of a neuron or an IPC, respectively) were restricted to deep layers of the cortex (Figures 2F–2M). However, in these experiments we also identified a small fraction of single-cell and two-cell clones in superficial layers of the neocortex (Figures 2F–2M). This suggested that some VZ progenitor cells generate pyramidal cells for superficial layers of the neocortex in their earliest neurogenic divisions.

We noticed that the clonal size of laminar-restricted lineages is typically smaller than that of translaminar clones (Figure 2E). One explanation for this difference could be that laminar-restricted lineages represent sub-clones resulting from the labelling of IPCs that undergo more than one round of cell division, generating four to five neurons with a laminar-restricted distribution. To test this hypothesis, we carried out lineage tracing experiments at single cell resolution using low-dose tamoxifen administration in *Tbr2^CreER^*;*RCE* pregnant mice at E12.5 (Figure S2A), which led to the sparse labeling of IPCs and their progenies (Pimeisl et al., 2013). We analyzed 73 IPC-derived lineages at P21 and exclusively found one-cell and two-cell clones, with no evidence for larger clones within our sample (Figures S2B–S2D). Although the existence of IPCs that undergo more than one cell division cannot be completely excluded, these results indicate that this is not common at this developmental stage. Consequently, IPCs are unlikely to be the origin of laminar-restricted lineages.

### Laminar-restricted lineages in the developing cortex

Retroviral tracing experiments suggested that the neurogenic output of neocortical VZ progenitor cells is significantly more heterogeneous than initially expected, including translaminar, deep- and superficial-layer restricted lineages. However, several technical limitations may contribute to the observation of laminar-restricted lineages, as retroviral tracing may lead to the incomplete labelling of neuronal lineages. For example, the existence of deep layer-restricted lineages might be due to the silencing of the viral cassette after a few rounds of cell division (Cepko et al., 2000), which would prevent the expression of GFP or mCherry in superficial layer PCs. There are also alternative explanations for the observation of superficial layer-restricted lineages in the retroviral tracing experiments that are independent from technical limitations. First, infected progenitors might have become neurogenic at slightly earlier stages and have already produced a wave of deep layer PCs before infection, which would therefore not be labeled by the retrovirus. Second, the entire set of deep layer neurons might have been generated during the first neurogenic division of a VZ progenitor cell, which would not be labeled in some cases due to the retroviral integration mechanism.

To overcome these technical limitations, we took advantage of the Mosaic Analysis with Double Markers (MADM) technique, a genetic method widely used to fate-map cellular lineages at high resolution (Hippenmeyer et al., 2010; Zong et al., 2005). We used the *Emx1-CreER^T2^* mice (Kessaris et al., 2006) to induce MADM sparse labelling of VZ progenitor cells following tamoxifen administration at E12.5 (Figure 3A). We specifically focused our analysis on G2-X MADM segregation events that result in the labelling of an unbalanced number of daughter cells with either green or red fluorescent proteins and report the outcome of asymmetric divisions in VZ progenitor cells (Zong et al., 2005). Consistent with the retroviral lineage tracing experiments, we found that the vast majority of MADM lineages adopt a translaminar configuration (Figures 3B and 3D). In addition, we also identified some lineages in which PCs were confined to layers V and VI, thereby confirming the existence of cortical lineages restricted to deep layers of the neocortex (Figures 2C and 2D). The fraction of deep layer-restricted lineages labelled with MADM (~7%) is smaller than that obtained with retroviral tracing (15%), which suggested that reporter silencing might exist in some clones in the retroviral lineage tracing experiments. In contrast, we did not recover a significant number of superficial layer-restricted lineages in MADM experiments (Figure 3D).

**Figure 3.**
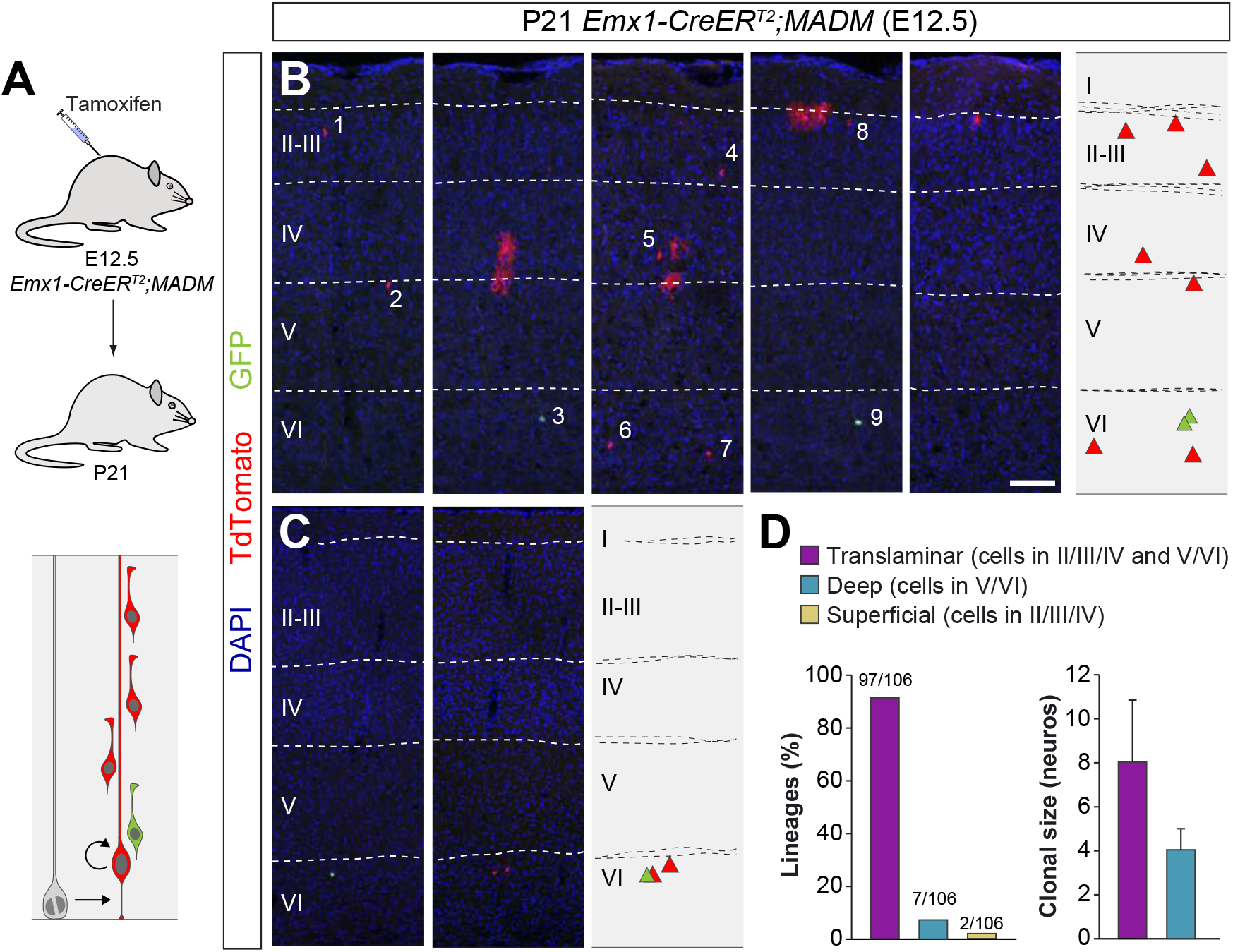
Lineage tracing using MADM identifies a small fraction of deep layer-restricted cortical lineages. (A) Experimental paradigm. The bottom panel illustrates the expected labeling outcome of a neurogenic RGC division following inducible MADM-based lineage tracing in which two subclones are labelled with different reporters. (B–C) Serial coronal sections through the cortex of P21 *Emx1-CreER^T2^;MADM* mice treated with tamoxifen at E12.5. The images show examples of translaminar (B) and deep layer-restricted (C) lineages. The schemas collapse lineages spanning across several sections into a single diagram. (D) Quantification of the fraction of translaminar, deep and superficial layer-restricted lineages, and clonal size in MADM lineages derived from a neurogenic (asymmetric) RGC division. Clonal size data are presented as mean ± standard deviation. Scale bar equals 100 μm.

The MADM experiments suggested that the observation of superficial layer-restricted lineages in retroviral experiments might be artifactual, a result of the incomplete retroviral labelling of neuronal lineages. We reasoned that if this were the case, the analysis of the MADM sub-clones (i.e., only one of the two colors in the lineage) containing more than two cells should lead to a similar fraction of ‘artifactual’ lineages, since these would essentially correspond to those labelled by retroviral infection missing the first division of VZ progenitor cells (Figure S3). This analysis indeed identified a small fraction of MADM sub-clones as ‘apparent’ superficial layer-restricted lineages (~12%), which was nevertheless significantly smaller than those identified in the retroviral experiments (22%). This indicated that although some of the superficial layer-restricted lineages observed in retroviral experiments were artifactual, others might not be.

One important difference between both approaches is that MADM G2-X recombination events occur exclusively in mitotic cells (Zong et al., 2005), while retroviral labelling does not strictly depend on cell division. Retroviruses require cell division for their integration into the genome, but the infection is independent of cell cycle stage (Cepko et al., 2000). Thus, we hypothesized that MADM may not consistently label a fraction of quiescent or slowly dividing progenitors, which could otherwise be targeted by retroviral infection. To test this idea, we carried out a new set of lineage tracing experiments using a third, complementary method. In brief, we traced cortical lineages at single cell resolution using low-dose tamoxifen administration in *Emx1-CreER^T2^;RCE* pregnant mice at E12.5 (Figure 4A), in which labelling of VZ progenitor cells should be independent of cell cycle dynamics. Since this method does not distinguish between lineages derived from symmetric or asymmetric cell divisions, we limited our analysis to lineages with a maximum of 12 cells, the larger clonal size of neurogenic lineages in the *Emx1-CreER^T2^;MADM* dataset (clones with more than 12 cells account for less than 5% of the neurogenic lineages and largely include [87%] the outcome of symmetrically-dividing progenitor cells).

**Figure 4.**
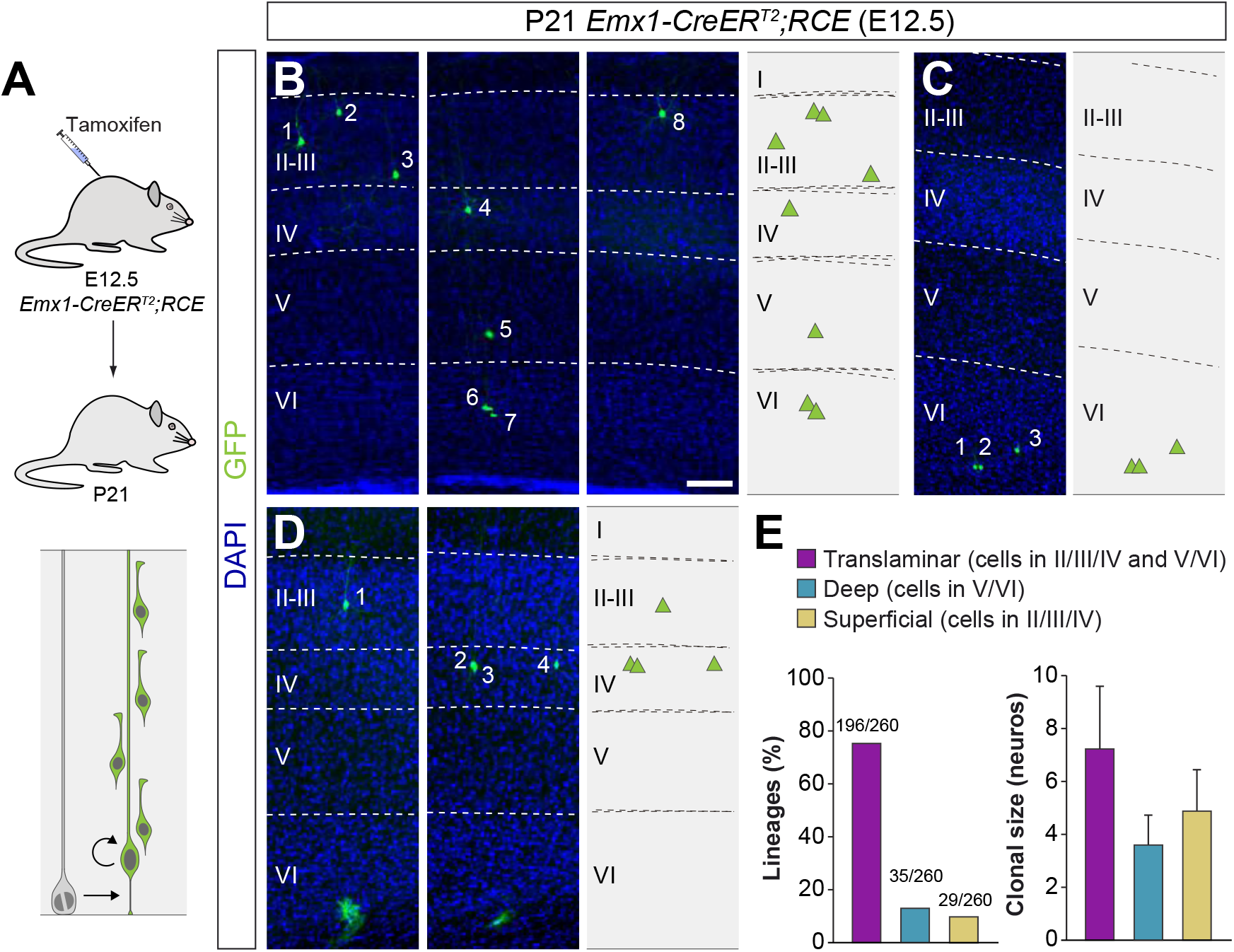
A fraction of early-quiescent cortical progenitors generates superficial layer-restricted lineages. (A) Experimental paradigm. The bottom panel illustrates the expected labeling outcome of a neurogenic RGC division following inducible conditional reporter lineage tracing in *Emx1-CreER^T2^;RCE* mice. (B–C) Serial coronal sections through the cortex of P21 *Emx1-CreER^T2^;RCE* mice treated with low-dose tamoxifen at E12.5. The images show examples of translaminar (B), deep layer-restricted (C) and superficial layer-restricted (D) lineages. The schemas collapse lineages spanning across several sections in a single diagram. (E) Quantification of the fraction of translaminar, deep and superficial layer-restricted lineages, and clonal size in inducible conditional reporter lineage tracing experiments. Clonal size data are presented as mean ± standard deviation. Scale bar equals 100 μm.

Consistent with the other approaches, the majority of lineages (~75%) labeled by injection of *Emx1-CreER^T2^;RCE* mice with low tamoxifen doses at E12.5 were translaminar (Figures 4B and 4E). We also confirmed that ~13% of the lineages were restricted to deep cortical layers (Figures 4C and 4E). In addition, we found that ~11% of the lineages consist of PCs confined to superficial layers of the neocortex (Figures 4D and 4E). In sum, the combined results of three different sets of lineage tracing experiments suggest that translaminar (~80%), deep layer-restricted (~10%) and superficial layer-restricted (~10%) lineages are generated at the onset of neurogenesis in the developing neocortex.

### Pyramidal cell lineages acquire diverse configurations

We next explored the precise organization of cortical lineages derived from VZ progenitor cells at E12.5. In lineage tracing experiments using *Emx1-CreER^T2^;RCE* mice (Figure 5A), we observed that only about a quarter of traced lineages contains neurons in every cortical layer from II to VI, and every other clone lacks neurons in one or multiple cortical layers (Figures 5B–D and 5F). For instance, a significant proportion of translaminar lineages lack PCs in layer V (Figures 5B and 5F) or layer IV (Figures 5C and 5F) but, considered collectively, PC lineages adopt every possible configuration of laminar distributions in the neocortex (Figure 5F). The heterogeneous organization of cortical lineages was not exclusively observed in the experiments performed in *Emx1-CreER^T2^;RCE* mice; similar results were obtained in retroviral lineage tracing and MADM experiments (Figure S4). Of note, the clonal size of translaminar lineages also exhibits heterogeneity (Figures 5E and S4).

**Figure 5.**
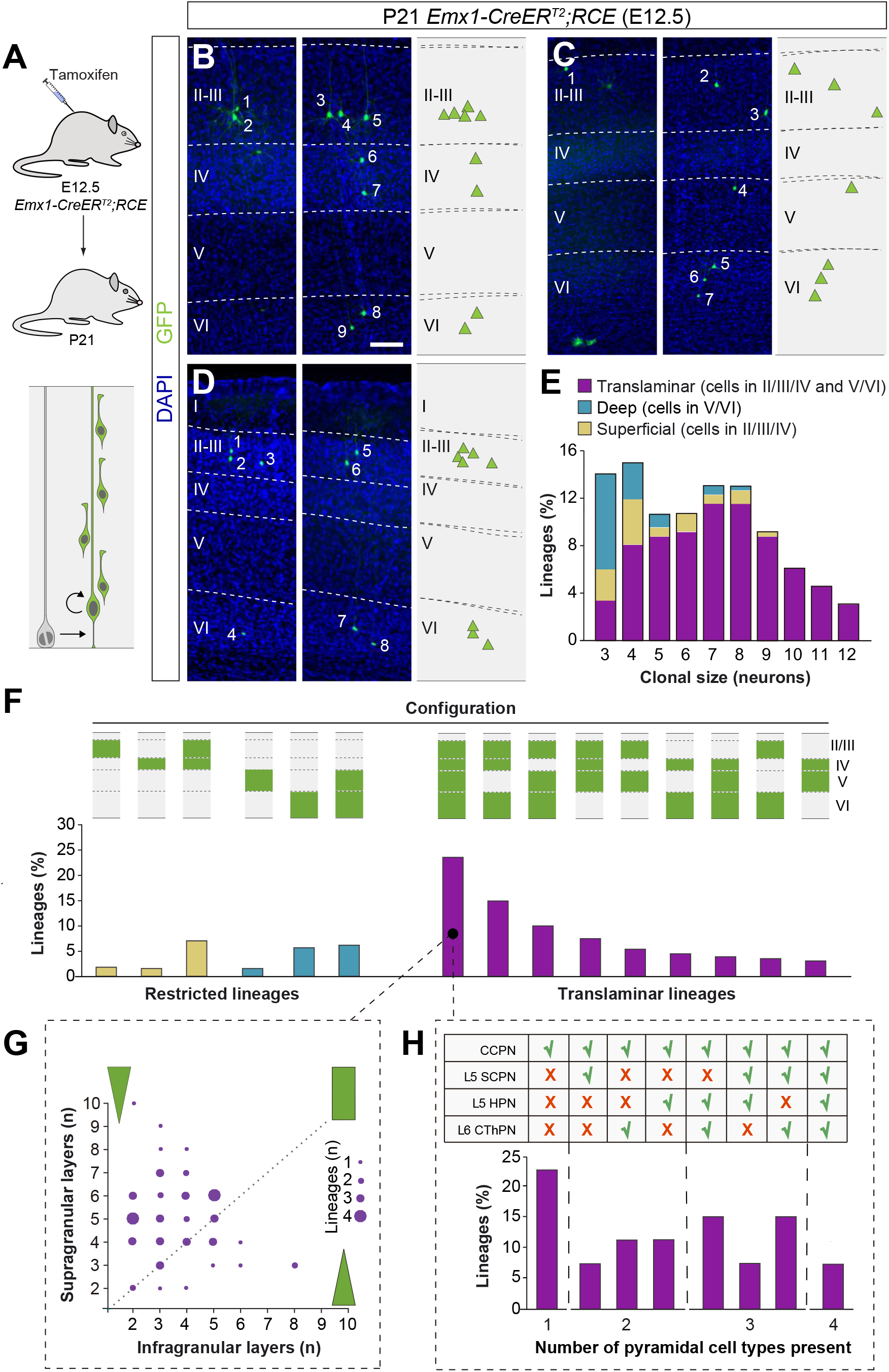
Translaminar lineages adopt very heterogeneous configurations. (A) Experimental paradigm. The bottom panel illustrates the expected labeling outcome of a neurogenic RGC division following inducible conditional reporter lineage tracing in *Emx1-CreER^T2^;RCE* mice. (B–xD) Serial coronal sections through the cortex of P21 *Emx1-CreER^T2^;RCE* mice treated with low-dose tamoxifen at E12.5. The images show examples of translaminar lineages with various laminar configurations. The schemas collapse lineages spanning across several sections into a single diagram. (E) Clonal size distribution of translaminar, deep and superficial layer-restricted lineages. (F) Relative frequency (expressed as percentage over the total number of lineages) of the different laminar configurations (green and grey schemas) in inducible conditional reporter lineage tracing experiments. (G) Relative abundance of PCs in superficial and deep layers from translaminar lineages containing cells in every layer. Lineages are represented as circles in a bi-dimensional space, indicating the number of cells in superficial versus deep layers. The size of the circle indicates the number of lineages that shown a particular configuration. Green shapes schematically represent lineage configurations. A rectangular shape illustrates lineages with a balanced number of superficial and deep PCs; triangular shapes represent configurations of lineages biased towards superficial or deep layer neurons. (H) Fraction of translaminar lineages with neurons in every layer containing one, two, three or four subclasses of PCs. CCPN, cortico-cortical projection neuron; SCPN, subcortical projection neuron; HPN, heterogeneous projection neuron; CThPN, cortico-thalamic projection neuron. Scale bar equals 100 μm.

We characterized the organization of translaminar lineages with PCs in every layer by quantifying the relative proportion of neurons in deep and superficial layers. This analysis revealed that these cortical lineages typically show a bias toward the production of PCs for superficial layers, although a minority of lineages displayed a preference toward deep layers identities or a balanced distribution across deep- and superficial layers (Figure 5G). To explore further the molecular diversity of PCs in these lineages, we stained P21 brain sections from *Emx1-CreER^T2^;RCE* mice induced at E12.5 with antibodies against Ctip2 and Satb2, two transcription factors whose relative expression defines different types of PCs with unique patterns of axonal projections (Greig et al., 2013; Lodato and Arlotta, 2015). We identified four PC subclasses based on the expression of these markers and their laminar distribution (Figure S5, also see Methods): Corticocortical projection neurons (CCPN), subcerebral projection neurons (SCPN), corticothalamic projection neurons (CThPN) and heterogeneous projection neurons (HPN) (Harb et al., 2016). Using this classification, we found that nearly a quarter of all-layer translaminar lineages were composed exclusively by CCPNs, while multiple different combinations of PC identities comprise the remaining lineages (Figure 5H). Of note, only a minor fraction of all cortical lineages contains the entire complement of subtypes identified. Altogether, our experiments reveal that PC lineages exhibit a great degree of heterogeneity in the number and identities they comprise.

### Heterogeneous lineage configurations arise from progenitor cells

The observed heterogeneity in cortical lineages likely emerges during neurogenesis. However, it is possible that selective cell death of specific PCs might contribute to the heterogeneous organization of cortical lineages. Recent studies have shown PCs undergo apoptosis during early postnatal stages (Blanquie et al., 2017; Wong et al., 2018). To explore the contribution of cell death to the heterogenous configuration of cortical lineages, we labeled clones by injecting a low dose of tamoxifen in *Emx1-CreER^T2^;RCE* mice at E12.5 and analyzed their laminar organization at P2, prior to the period of PC death (Wong et al., 2018). We detected no significant differences in the average clonal size or in the relative frequency of P2 translaminar and laminar-restricted lineages compared to P21 (Figures S6A–S6D). In addition, we observed that the diversity of lineage patterns was remarkably similar between P2 and P21 (Figure S6E). We also noticed a tendency (*χ*^2^ test, *p* = 0.099) for the fraction of lineages with PCs in every layer to be overrepresented and the frequency of lineages lacking PCs in layer 5 to be underrepresented at P2 (Figures S6E). These experiments revealed that although cell death may have a subtle impact in refining the final diversity of lineages and their relative proportion, it is likely that such heterogeneity arises directly during the process of cortical neurogenesis.

### A small number of progenitor identities underlies lineage diversity

The variability in size and composition of PC lineages raises questions about the diversity of VZ progenitor cells. One possibility to explain such heterogeneity in lineage configurations is the existence of equipotent VZ progenitor cells that are subject to stochastic factors controlling lineage progression, as proposed for the retina (He et al., 2012). Alternatively, the diversity of lineage configurations might be due to different types of VZ progenitor cells with restricted potential to generate specific classes of PCs (Franco and Muller, 2013).

We used a mathematical approach based on *in silico* simulations to identify the most likely model explaining the existence of heterogeneous PC lineages. To avoid variability that could be attributed to the distribution of lineages across different areas of the neocortex, experimental data was exclusively derived from lineages mapped in the primary somatosensory cortex (S1) of *Emx1-CreER^T2^;RCE* mice. We define a series of control outcomes, based on our experimental observations, which any model should replicate to be considered valid. First, the relative laminar proportion of PCs generated by modelled lineages should be consistent with the experimental results. Second, modeled lineages should reproduce the clonal size distribution observed in our experiments. Finally, experimental lineages exhibit a negative correlation between the number of PCs in deep and superficial layers; consequently, as a third condition, modelled lineages should collectively exhibit a similar correlation. We calculated a z-score that represents the distance between the simulated dataset and the experimental data for each of these parameters; models were considered valid only if the z-score for all three parameters was < 1 (a z-score of zero would represent no difference between simulated and experimental data; see Methods for details).

Before considering the possible existence of one or more types of progenitor cells, we wondered whether the heterogeneous organization of cortical lineages has any actual impact on the structure of the cortex. In other words, we asked whether lineage structure is indeed relevant for laminar organization. It is formally possible that the diversity of cortical lineages is simply the consequence of a random process on PC generation in which the only boundary condition is the relative number of PCs that populate each layer of the cortex. To test this hypothesis, we randomly permuted the PCs obtained from the different lineages while maintaining their laminar identities and the total number of cells in each lineage. If this model were correct, permuted lineages should reproduce experimental data. As expected, this process reproduced the clonal size distribution and number of cells per layer observed in the experimental data (Figures S7A and S7B). However, it failed to replicate the observed anti-correlation in neuron numbers that exists between superficial and deep layers (Figures S7C). In addition, we observed that the laminar configuration of the permuted lineages differed from the experimental data (data not shown). These results indicated that the laminar distribution of neurons in each lineage arises specifically from an organized pattern of neurogenesis.

We next tested whether a unique equipotent population of VZ progenitor cells, with capacity to perform a number of stochastic decisions, could generate the observed diversity of cortical lineages. To test this hypothesis, we modelled cortical progenitor behavior using three basic rules (Figure S7D). First, *in silico* progenitors would generate neurons of different layers sequentially, following the experimentally observed inside-out pattern. Second, *in silico* progenitors would have a number of opportunities to generate neurons in each layer that corresponds to the maximum number of neurons found for each particular layer in a single lineage among our three experimental datasets. Third, the decision to generate a neuron would be probabilistic, with cell generation probabilities varying by cortical layer but equal for all opportunities within the same layer. Using this set of rules, we then systematically explored combinations of generation probabilities in an effort to replicate the lineage configurations observed experimentally. We found that a model with a single set of generation probabilities (i.e., an equipotent population of VZ progenitor cells: Model 1) could not satisfy all control parameters simultaneously. For instance, when modeled lineages reproduced the laminar organization of the cortex and the observed superficial-deep anti-correlation in neuronal numbers (Figures S7E and S7G), they failed to reproduce the experimental clonal size distribution (Figure S7F).

We then introduced a second population of progenitor cells with a different set of probabilities for cell generation (Model 2, Figure 6A). In this model, we found a set of generation probability values for which lineages satisfied all three control parameters (Figures 6B–6D). In addition, we observed that the relative proportions of translaminar and laminar-restricted lineages were identical to those measured experimentally (Figure 6E). Finally, translaminar modeled lineages showed similar laminar configurations when compared to lineages obtained experimentally (Figure 6F). In sum, mathematical modelling suggests that two different progenitor cell populations, each generating neurons sequentially with a different stochastic rule, are sufficient to generate the observed diversity of cortical lineage distributions.

**Figure 6.**
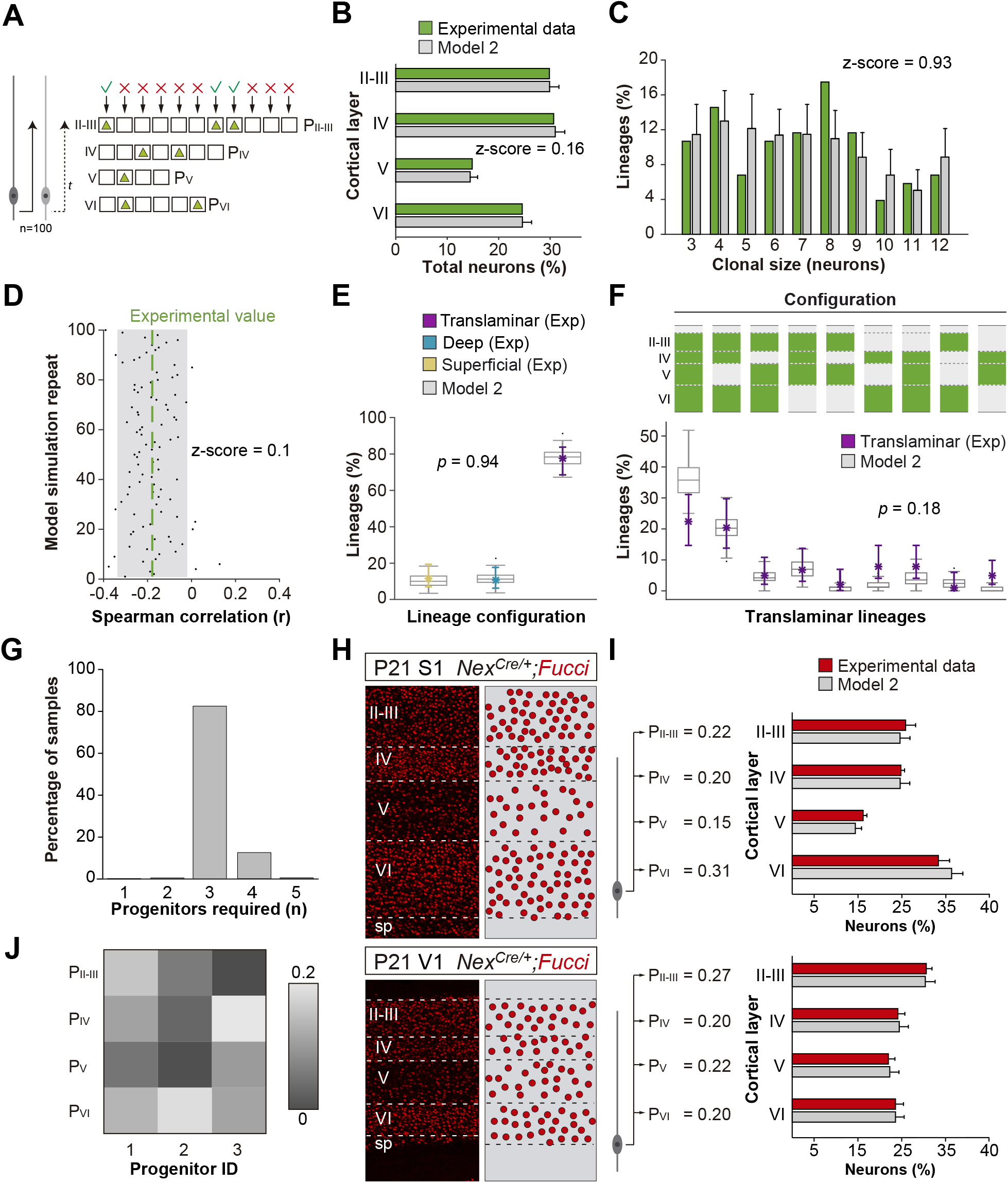
A small number of progenitor identities can generate the diversity of cortical lineages. (A) Schematic representation of a mathematical model of cortical neurogenesis in which two different progenitor identities are modeled (Model 2). Squares represent the number of stochastic decisions performed by each progenitor for each cortical layer during in silico simulations. The odds of generating neurons for each chance are given by a probability value (P), which is unique for each layer and progenitor identity. The model runs 100 simulations with 100 progenitors. (B) Fraction of cells in each cortical layer (expressed as percentage of total) in experimental and modeled lineages. (C) Clonal size distribution in experimental and modeled lineages. (D) Spearman correlation (r) values for the fraction of superficial and deep layer neurons in modeled lineages. Each dot represents an r value for one simulation. The green line shows the experimental value; the shadow area around the experimental data represents a 95% confidence interval for the experimental value. (E) Fraction of translaminar, deep and superficial layer-restricted lineages found experimentally and predicted by the model (expressed as percentage of all modeled lineages within a single simulation). Grey boxes represent variability among 100 simulations; colored stars and lines show experimental values and 95% confidence intervals for experimental values (*p* = 0.94, Fisher’s exact test). (F) Relative frequency (expressed as percentage over all modeled translaminar lineages within a single simulation) of laminar configurations in experimental and modeled translaminar lineages. Grey boxes represent variability among 100 simulations; colored stars and lines show experimental values and 95% confidence intervals for experimental values (*p* = 0.18, Chi-square test). (G) Distribution of the number of progenitor identities required to describe the data using Bayesian modeling. Posterior probabilities were obtained from 2000 posterior samples of the model parameters (see Methods). (J) Occupancy probability values obtained from the Bayesian model in the case of three different progenitor identities. Numerical estimates correspond to posterior averages across 2000 samples. (H) Coronal sections through the primary somatosensory (S1) and visual (V1) cortex of P21 *Nex^Cre/+^;Fucci* mice. The schemas on the right illustrate PC densities per layer. (I) Fraction of PCs per layer (expressed as percentage of total neurons) generated with two sets of laminar probability factors using Model 2 compared to the experimental data. Histograms represent mean ± standard deviation. Z-scores represent the distance between experimental and simulated datasets for each parameter, which is calculated as the difference between the averages of model and experimental data divided by the standard deviation within model simulations (see Methods).

We noticed, however, that modeled lineages did not achieve perfect reproduction of the experimental data. This may suggest that, while a minimum of two progenitor cell populations could in essence replicate the diversity of some experimental configurations, the generation of such diversity in vivo may generally involve additional progenitor identities. To explore this idea, we made use of Bayesian modeling to infer the probability distribution of the number of progenitor identities conditional to the lineages observed experimentally across all possible model architectures (see Methods for details). Consistent with the previous results, Bayesian inference on the number of progenitor identities revealed that a single progenitor cell population is very unlikely to reproduce the observed diversity in cortical lineages (Figure 6G). However, the posterior distribution of the number of progenitor identities has a sharp peak (85%) corresponding to three different progenitor cell populations to describe the data (Figures 6G, 6J and S8). These results indicate that a small number of progenitor cell identities following different stochastic developmental programs likely underlies the generation of the diversity of PC lineage configurations.

Finally, we explored whether the proposed stochastic model, which involves at least two types of progenitor cells, would be able to generate different ratios of layer-specific neurons under different circumstances (i.e., different cell generation probabilities). This would robustly account for the emergence of cytoarchitectural differences across neocortical areas. To this end, we quantified the fraction of PCs in each layer of the primary somatosensory (S1) and visual (V1) cortices in *Nex-Cre;Fucci2* mice, in which all pyramidal neurons in the neocortex are labeled with a nuclear fluorescent marker. As expected, we found important differences in laminar cytoarchitecture between both regions (Figure 6H). Remarkably, we found that subtle tuning of division probabilities for both areas replicated the different laminar ratios in our model (Figure 6I). This suggests that two or perhaps three cortical VZ progenitor types, following independent stochastic behavioral rules, would suffice to generate very diverse cytoarchitectonical patterns.

## DISCUSSION

In this study, we have analyzed the organization of hundreds of pyramidal cell lineages using three complementary methods. Our results indicate that the output of individual progenitor cells in the developing mouse neocortex is much more heterogeneous than previously anticipated. The most abundant progenitor cells in the pallium generate pyramidal cells for both deep and superficial layers of the cortex, as suggested by previous studies. Unexpectedly, however, a sizable fraction of those lineages lacks pyramidal cells in one or several layers. The heterogeneous output of cortical progenitor cells include lineages in which PCs are restricted to either deep or superficial layers. Mathematical modeling suggests that a minimum of two progenitor cell populations, each generating PCs sequentially with a different stochastic rule, could generate the diversity of individual cortical lineage configurations from which the complex cytoarchitecture of the neocortex emerges.

### Methodological considerations

Understanding how individual lineages contribute to the production and organization of pyramidal cells is essential to articulate a coherent framework of cortical development. The analysis of the output of progenitor cells in the developing rodent cortex expands over three decades and has relied on four approaches: retroviral labeling (Luskin et al., 1988; Noctor et al., 2001; Noctor et al., 2004; Price and Thurlow, 1988; Reid et al., 1995; Walsh and Cepko, 1988, 1992), mouse chimeras (Tan et al., 1998), MADM (Beattie et al., 2017; Gao et al., 2014) and genetic fate-mapping (Eckler et al., 2015; Franco et al., 2012; Garcia-Moreno and Molnar, 2015; Gil-Sanz et al., 2015; Guo et al., 2013; Kaplan et al., 2017). These studies often led to contradictory results, which has prevented the emergence of a consistent model. The prevalent view is that each progenitor cell in the developing pallium is multipotent and generates a cohort of pyramidal cells that populate all layers of the neocortex except layer I (Eckler et al., 2015; Gao et al., 2014; Guo et al., 2013; Kaplan et al., 2017), as originally conceived in the “radial unit hypothesis” (Rakic, 1988). In contrast, some authors have suggested that many cortical progenitor cells are fate-restricted to generate pyramidal cells that exclusively occupy deep or superficial layers of the neocortex (Franco et al., 2012; Franco and Muller, 2013; Gil-Sanz et al., 2015). Here, we have used three different methods (retroviral labeling, MADM and genetic fate-mapping) to investigate the clonal production of cortical neurons by capitalizing on the synergy that emerges from the advantages of each individual approach. Our results indicate that this multi-modal approach is required to comprehensively capture the complex behavior of progenitor cells in the developing cortex.

Retroviral labeling has two important limitations: it only labels hemi-lineages and is prone to silencing, which may prevent the identification of the entire progeny of a progenitor cell (Cepko et al., 2000). Conversely, retroviral labeling targets progenitor cells indiscriminately and, consequently, is not biased towards a particular genetic fate (Cepko et al., 2000), as is the case for genetic strategies. MADM, on the other hand, has the enormous advantage of identifying both sister cells resulting from a cell division.

However, G2-X MADM events require progenitor cells to undergo cell division at the time of induction because it directly relies on Cre-dependent inter-chromosomal mitotic recombination (Zong et al., 2005). Our results revealed that MADM does not reliably label a small fraction of progenitor cells present in the pallial VZ at E12.5 that gives rise to cohorts of pyramidal cells exclusively located in superficial layers of the neocortex. These lineages were however observed in both retroviral labeling experiments and in genetic tracing experiments using the same genetic driver (*Emx1-CreER^T2^*) as in the MADM experiments, which strongly suggests that some Emx1+ progenitor cells producing exclusively superficial layer PCs in the developing cortex are not targeted by the MADM approach. We hypothesize that these progenitors might be quiescent or slow-dividing progenitors at this stage and become more active at later stages of development. Finally, although the use of genetic fate-mapping strategies (e.g., *Emx1-CreER^T2^;RCE*) is a powerful method to investigate cortical lineages, it has the important constraint of not being able to distinguish between symmetric proliferating and asymmetric neurogenic divisions. This hampers the analysis of clonal sizes, which can be otherwise accurately assessed with MADM except for the lineages that are not detected with this method.

### Diversity of neocortical lineages

Previous clonal analyses based on MADM lineage tracing experiments led to the suggestion that individual progenitor cells in the pallial VZ produce a unitary output of approximately 8 excitatory neocortical neurons (Gao et al., 2014). However, these experiments also revealed a high degree of variability of clonal sizes (Gao et al., 2014). In agreement with this notion, our experiments reveal that individual cortical progenitors generate variable numbers of neurons, typically ranging from 3 to 12, with some exceptional outliers. Moreover, as previous MADM studies did not reveal lineages with restricted laminar patterns (either deep or superficial layer restricted clones), they underestimated the fraction of lineages with relatively small clonal size. Consequently, our analysis of neurogenic lineages revealed a bimodal distribution of clonal sizes with defined peaks centered at 4 and 8 cells, which largely correspond to the contribution of laminar-restricted and translaminar lineages, respectively (Figure 5E).

Our study also revealed that individual cortical progenitor cells generate lineages with very diverse combinations of pyramidal cell types. Cortical progenitors are thought to undergo progressive changes in their competency to generate different layer-specific types of pyramidal neurons (Desai and McConnell, 2000; Rakic, 1974). Consistent with this idea, our results reveal that most cortical progenitors generate diverse types of excitatory neurons. However, since many cortical progenitor cells fail to generate neurons for at least one layer of the neocortex, the majority of cortical lineages does not include the diversity of excitatory neurons. In other words, the fraction of individual cortical lineages that would be considered as “canonical” – i.e., containing all three main classes of excitatory projection neurons (CCPN, SCPN and CThPN) – is significantly smaller than previously anticipated. Considering the variance in clonal size and lineage composition of neocortical lineages, our results indicate that cortical progenitor cells exhibit very heterogeneous patterns of neuronal generation and specification.

### Fate-restricted neocortical progenitor cells

It has been suggested that some neocortical progenitor cells in the developing pallium generate pyramidal cells that are restricted to certain fates and layers of the neocortex (Franco et al., 2012; Gil-Sanz et al., 2015). In our experiments, approximately one in six cortical progenitor cells generate laminar-restricted lineages. The existence of lineages restricted to deep layers of the neocortex was observed with all three methods used in this study. Although some variation exists in the relative fraction of deep layer-restricted lineages observed with the different approaches, these differences lie within the expected experimental noise considering the relatively small number of lineages that belong to this category. In addition, both retroviral labelling and genetic fate-mapping experiments identified a fraction of cortical progenitor cells that generate pyramidal cells that exclusively populate the superficial layers of the neocortex. The discrepancy between the results of genetic fate-mapping and MADM experiments, in which the same mouse strain is used as the driver for recombination (*Emx1-CreER^T2^*), suggests that these fate-restricted lineages arise from progenitor cells that are not actively dividing at E12.5. This hypothesis is consistent with the identification of a population of self-renewing progenitors with limited neurogenic potential during the earliest phases of corticogenesis (Garcia-Moreno and Molnar, 2015).

The existence of superficial layer-restricted cortical lineages is further supported by the identification of intermediate progenitor cells as early as E12.5 that generate superficial layer excitatory neurons (this study and Mihalas et al., 2016), when the majority of deep layer pyramidal cells are being generated. Since intermediate progenitor cells derive from VZ progenitor cells (Haubensak et al., 2004; Miyata et al., 2004; Noctor et al., 2004) and cortical neurogenesis begins at these stages in the mouse (this study and Guo et al., 2014), this observation reinforces the idea that some cortical progenitors are fate-restricted to generate superficial layer pyramidal cells from early stages of corticogenesis.

It is presently unclear whether lineages with neurons restricted to deep or superficial layers of the neocortex arise from distinct pools of progenitor cells or should simply be considered as extreme examples of the enormous diversity of lineage configurations uncovered by our study. Previous studies have identified morphological heterogeneity among pallial VZ progenitor cells (Gal et al., 2006). However, there is limited evidence for important molecular differences among these cells (Mizutani et al., 2007; Pollen et al., 2014; Telley et al., 2016). In the absence of a definitive molecular signature, our results using mathematical modelling suggest that it is very unlikely that a homogeneous population of progenitor cells following a common developmental program could generate the observed diversity of neocortical lineages. Rather, a small number (two or more) of developmental programs (i.e., progenitor identities) likely underlies such diversity.

### A stochastic model of cortical neurogenesis

In spite of the great diversity of configurations that exist among individual neocortical lineages, our results suggest that a small number of progenitor cell identities (2 or 3) with distinct developmental programs would be able to generate the whole diversity of observed outcomes. Our model assumes that progenitor cells have the potential to generate any type of excitatory neuron subtype, but that distinct “types” of progenitor cells would have different probabilities to generate pyramidal cells during development. Using such a model, a single progenitor cell type – a “canonical” radial glia cell – is able to reproduce a great diversity of lineage configurations (essentially all of the translaminar configurations), but strongly underestimate deep and superficial layer-restricted linages, which suggests that they likely arise from a different pool of progenitor(s). The model proposed here is somewhat reminiscent to that described for the developing zebrafish retina (He et al., 2012). In contrast to our findings, however, the entire diversity of lineages found in the zebrafish retina seem to be recapitulated by a single type of equipotent progenitor cells subjected to stochastic rules.

Cell death likely contributes to shaping the heterogeneous behavior of progenitor cells in the developing cortex. However, our data indicate that the configuration of cortical lineages prior to the normal period of programed cell death of pyramidal cells (Blanquie et al., 2017; Wong et al., 2018) is not significantly different that in juvenile mice, which suggests that cell death may shape but does not create the heterogeneous configurations adopted by cortical lineages. It remains to be investigated how embryonic cell death, most notably of neurogenic progenitor cells, may impact the organization of pyramidal cell lineages (Mihalas and Hevner, 2018). However, it has been shown that apoptotic nuclei are very rare in the VZ during neocortical development (Haydar et al., 1999; Thomaidou et al., 1997).

Our study suggests that progenitor cells in different cortical areas are likely constrained by different probabilistic rules, which would contribute to the generation of the diverse cytoarchitectonic patterns that are found across the neocortex. How stochastic neurogenic decisions are made remains to be elucidated, but they likely depend on the influence of extrinsic and intrinsic signals on parameters such as cell cycle length, the asymmetric inheritance of cell components and the membrane potential of progenitor cells (Haydar et al., 2003; Lange et al., 2009; Pilaz et al., 2009; Roccio et al., 2013; Vitali et al., 2018; Wang et al., 2009). Local signals in different neocortical areas would contribute to the tuning of progenitor cell behaviors to output different cytoarchitectures without the requirement of regional-specific progenitor populations. Consequently, this model allows great flexibility in the generation of heterogeneous cortical cytoarchitectures without the requirement of a large number of progenitor identities. The specification of a small number of progenitor cells with competence to adapt their neurogenic behavior to different probabilistic rules based on their location within the neocortical neuroepithelium represents the simplest and most robust mechanism for the generation of cortical circuitry.

## Supporting information

## AUTHOR CONTRIBUTIONS

A.L., G.C. and O.M. conceived and developed the project. A.L. performed most wet-lab experiments and analyses with the support of E.S. and M.F-O., with the exception of the first series of retroviral labeling experiments which were performed by G.C. S.H. designed MADM experiments. R.B. analyzed MADM data generated with the support of C.S. A.L. and F.K.W. analyzed cell death in cortical lineages. A.L., M. Maravall and G.D. performed mathematical analyses. M. Meyer and S.J.A. provided essential reagents. A.L. and O.M. wrote the paper with input from all authors.

## ACKNOWLEDGEMENTS

We thank I. Andrew and S.E. Bae for excellent technical assistance, F. Gage for plasmids, and K. Nave (*Nex-Cre*) for mouse colonies. We thank members of the Marín and Rico laboratories for stimulating discussions and ideas. Our research on this topic is supported by grants from the European Research Council (ERC-2017-AdG 787355 to O.M and ERC-2016-CoG 725780 to S.H.) and Wellcome Trust (103714MA) to O.M. L.L. was the recipient of an EMBO long-term postdoctoral fellowship, R.B. received support from FWF Lise-Meitner program (M 2416) and F.K.W. was supported by an EMBO postdoctoral fellowship and is currently a Marie Skłodowska-Curie Fellow from the European Commission under the H2020 Programme.

