## Supplementary material for "Heterogeneous progenitor cell behaviors underlie the assembly of neocortical cytoarchitecture"

Supplementary Figures S1-S8

Supplementary Table S1

Methods

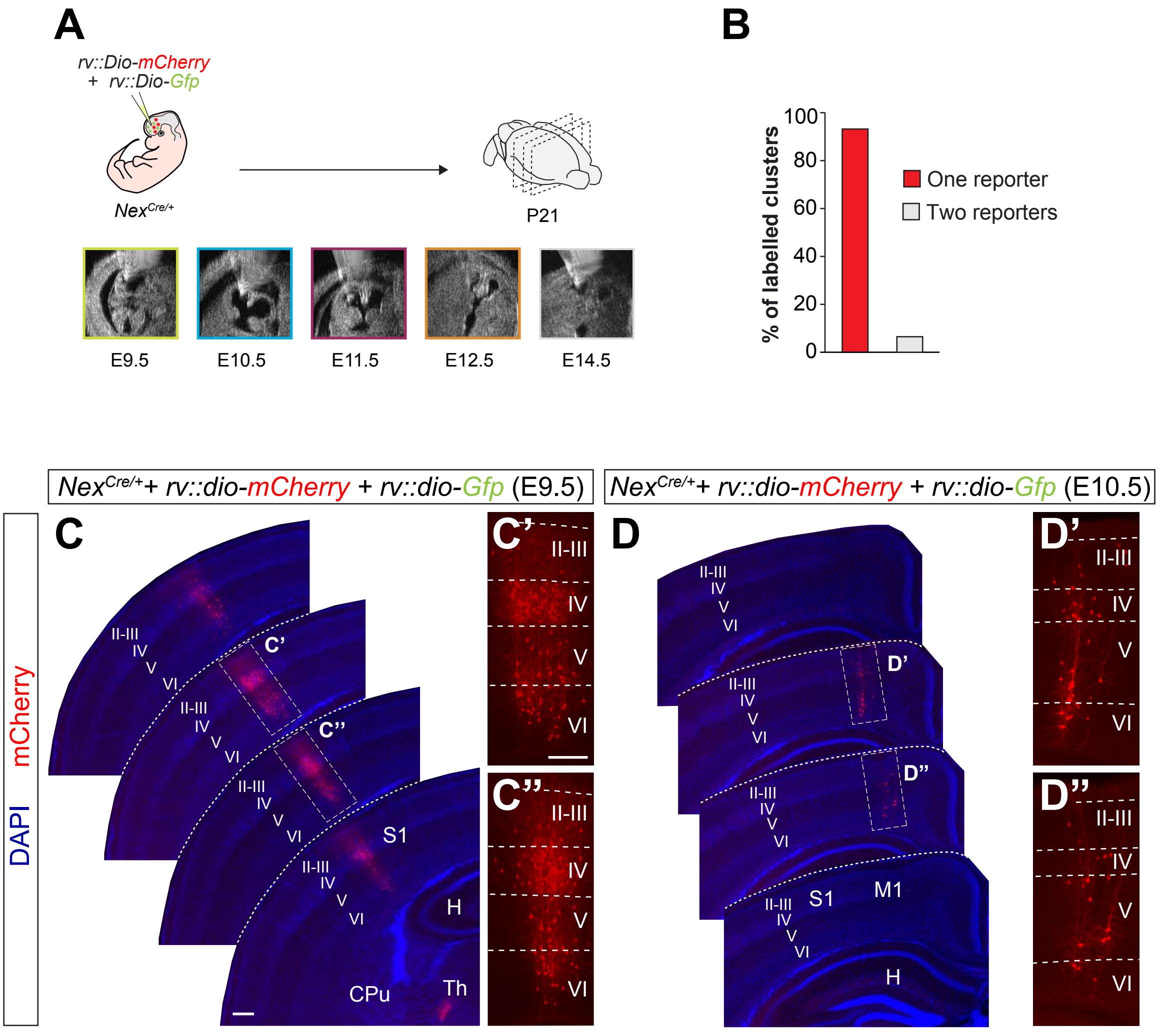
A
rv::Dio-mCherry
+ rv::Dio-Gfp
Nex^Cre/+
P21
E9.5
E10.5
E11.5
E12.5
E14.5
B
% of labelled clusters
100
80
60
40
20
0
One reporter
Two reporters
Nex^Cre/+ + rv::dio-mCherry + rv::dio-Gfp (E9.5)
Nex^Cre/+ + rv::dio-mCherry + rv::dio-Gfp (E10.5)
mCherry
DAPI
C
C'
C''
II-III
IV
V
VI
S1
H
CPu
Th
D
D'
D''
M1

**Figure S1. Sparse labeling of neuronal clones with low-titer retroviral infection**

(A) Experimental paradigm.
(B) Fraction of neuron clusters containing cells labeled with one or two reporters.
(C and D) Serial coronal sections through the telencephalon of P21 $Nex^{Cre/+}$ mice infected with low-titer conditional reporter retroviruses at E9.5 (C) and E10.5 (D). The high magnification pictures shown in (C' and C") and (D' and D") correspond to the clones shown in (C) and (D), respectively.
Scale bars equal 100 µm (C and D) and 300 µm (C', C", D' and D").

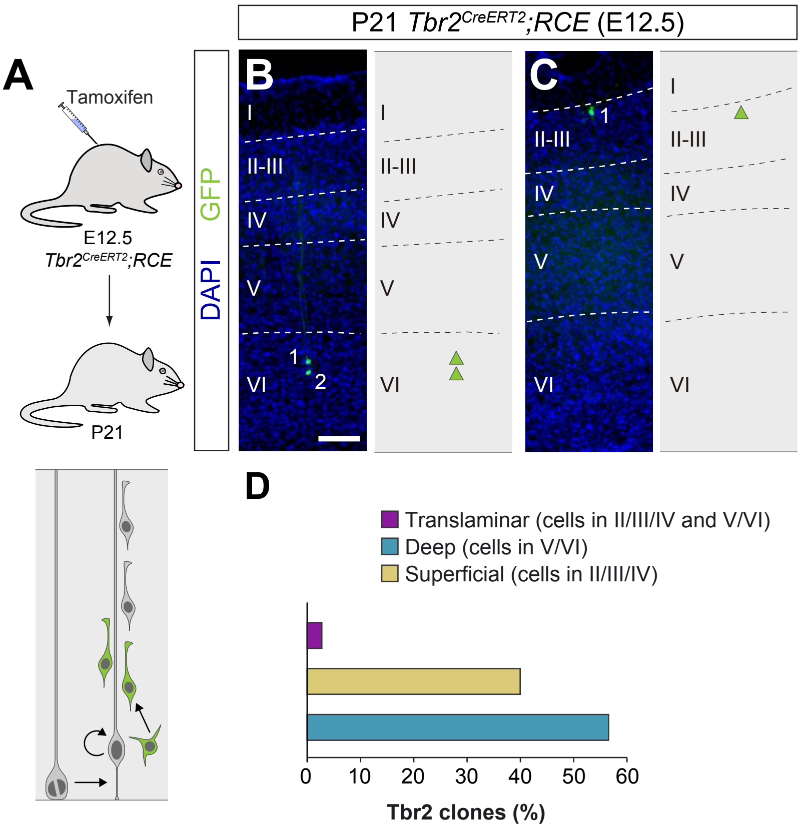
P21 Tbr2CreERT2;RCE (E12.5)
A
Tamoxifen
E12.5
Tbr2CreERT2;RCE
P21
B
C
GFP
DAPI
I
II-III
IV
V
VI
1
2
D
Translaminar (cells in II/III/IV and V/VI)
Deep (cells in V/VI)
Superficial (cells in II/III/IV)
0
10
20
30
40
50
60
Tbr2 clones (%)

**Figure S2. Lineage tracing of Tbr2+ intermediate progenitor cells**

(A) Experimental paradigm. The bottom panel illustrates the expected labeling outcome of the division of a Tbr2+ intermediate progenitor cell.

(B–C) Coronal sections through the cortex of P21 *Tbr2*$^{CreERT2}$*;RCE* mice treated with low-dose tamoxifen at E12.5. The images show examples of deep and superficial layer-restricted Tbr2-derived lineages. The schemas collapse lineages spanning across several sections into a single diagram.

(D) Laminar allocation of translaminar, deep and superficial layer-restricted Tbr2-derived lineages.

Scale bar equals 100 μm.

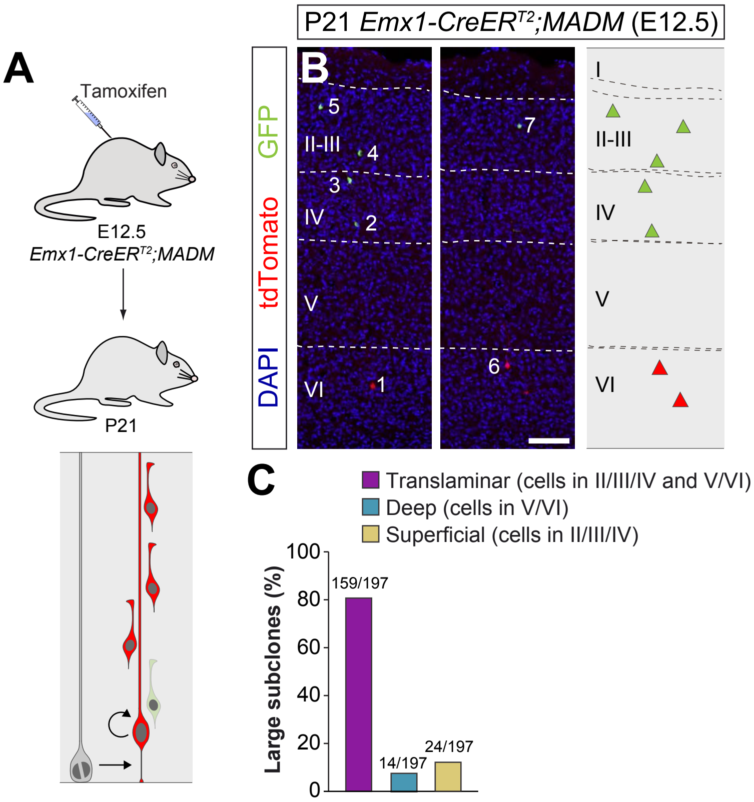
P21 Emx1-CreERT2;MADM (E12.5)
A
Tamoxifen
E12.5
Emx1-CreERT2;MADM
P21
B
GFP
tdTomato
DAPI
I
II-III
IV
V
VI
1
2
3
4
5
6
7
C
Translaminar (cells in II/III/IV and V/VI)
Deep (cells in V/VI)
Superficial (cells in II/III/IV)
Large subclones (%)
100
80
60
40
20
0
159/197
14/197
24/197

**Figure S3. Large MADM subclones reveal a small fraction of artifactual superficial layer-restricted in the retroviral dataset**

(A) Experimental paradigm. The bottom panel illustrates the expected labeling outcome of a neurogenic RGC division following inducible MADM-based lineage tracing in which two subclones are labelled with different reporters.
(B) Serial coronal sections through the cortex of P21 *Emx1-CreER*$^{T2}$*;MADM* mice treated with tamoxifen at E12.5. The images show an example of a translaminar lineage containing a large superficial layer-restricted sub-clone. The schemas collapse lineages spanning across several sections into a single diagram.
(C) Laminar configuration of large MADM sub-clones.
Scale bar equals 100 µm.

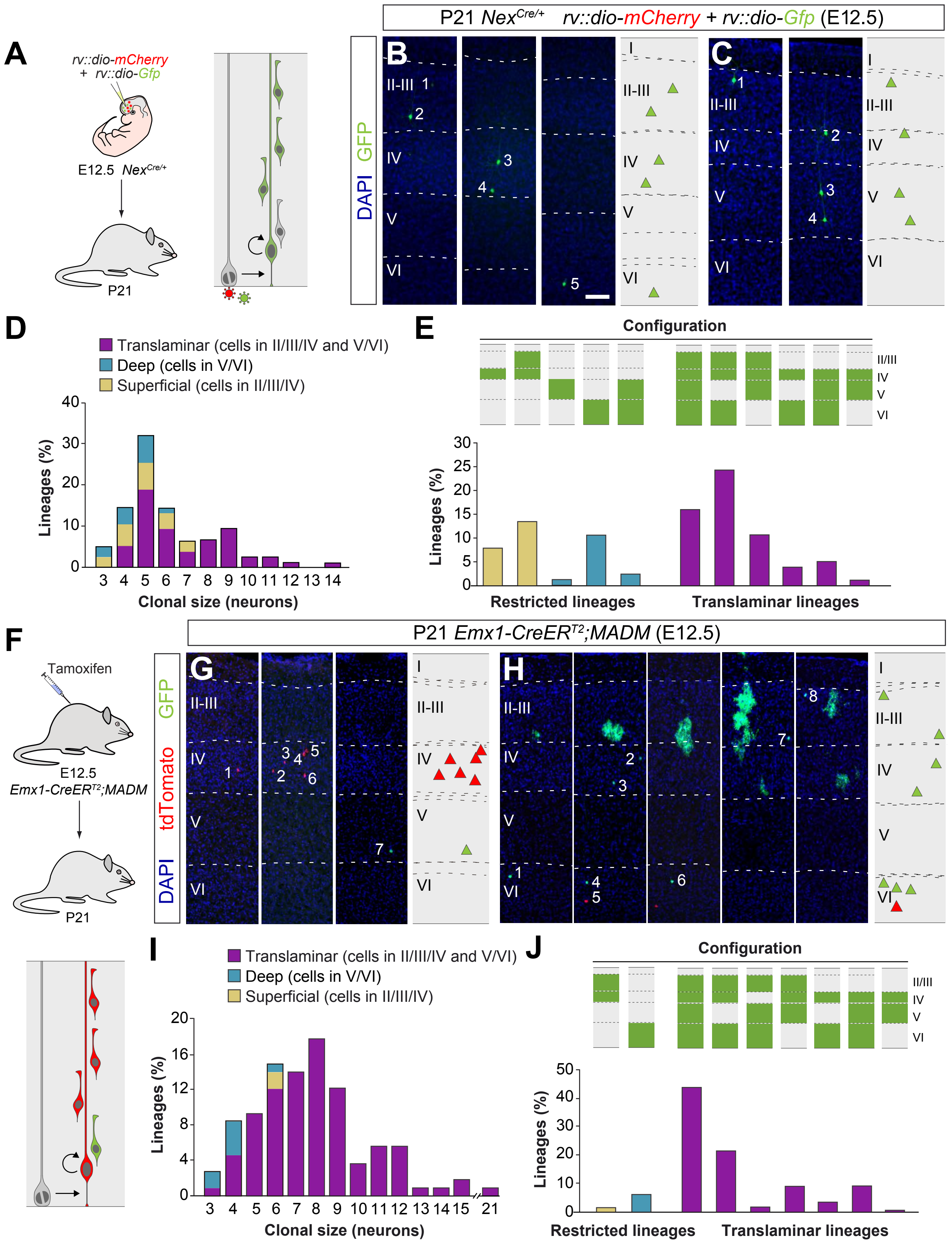
P21 Nex^Cre/+ rv::dio-mCherry + rv::dio-Gfp (E12.5)
A
rv::dio-mCherry
+ rv::dio-Gfp
E12.5 Nex^Cre/+
P21
B
C
DAPI GFP
II-III
IV
V
VI
I
D
Translaminar (cells in II/III/IV and V/VI)
Deep (cells in V/VI)
Superficial (cells in II/III/IV)
Lineages (%)
Clonal size (neurons)
E
Configuration
II/III
IV
V
VI
Lineages (%)
Restricted lineages
Translaminar lineages
F
P21 Emx1-CreER^T2;MADM (E12.5)
Tamoxifen
E12.5
Emx1-CreER^T2;MADM
P21
G
H
DAPI tdTomato GFP
I
Translaminar (cells in II/III/IV and V/VI)
Deep (cells in V/VI)
Superficial (cells in II/III/IV)
Lineages (%)
Clonal size (neurons)
J
Configuration
II/III
IV
V
VI
Lineages (%)
Restricted lineages
Translaminar lineages

**Figure S4. Retrovirus and MADM labelled lineages reproduce laminar configuration diversity**

(A) Experimental paradigm. The right panel illustrates the expected labeling outcome following retroviral infection of an RGC undergoing a neurogenic cell division in which the viral integration occurs in the self-renewing RGC.
(B–C) Serial coronal sections through the cortex of P21 $Nex^{Cre/+}$ mice infected with low-titer conditional reporter retroviruses at E12.5. The images show examples of layer V (B) and layer 6 (C) skipping lineages. Dashed lines define cortical layers. The schemas collapse lineages spanning across several sections into a single diagram.
(D) Clonal size distribution of translaminar, deep and superficial layer-restricted lineages in retroviral labeling experiments.
(E) Relative frequency (expressed as percentage over the total number of lineages) of the different laminar configurations (green and grey schemas) in retroviral labeling experiments.
(F) Experimental paradigm. The bottom panel illustrates the expected labeling outcome of a neurogenic RGC division following inducible MADM-based lineage tracing in which two subclones are labelled with different reporters.
(G–H) Serial coronal sections through the cortex of P21 $Emx1\text{-}CreER^{T2};MADM$ mice treated with tamoxifen at E12.5. The images show examples of layer IV and V restricted (G) and layer V skipping (H) lineages. The schemas collapse lineages spanning across several sections into a single diagram.
(I) Clonal size distribution of translaminar, deep and superficial layer-restricted lineages in MADM labeling experiments.
(J) Relative frequency (expressed as percentage over the total number of lineages) of the different laminar configurations (green and grey schemas) in MADM labeling experiments.
Scale bar equals 100 µm.

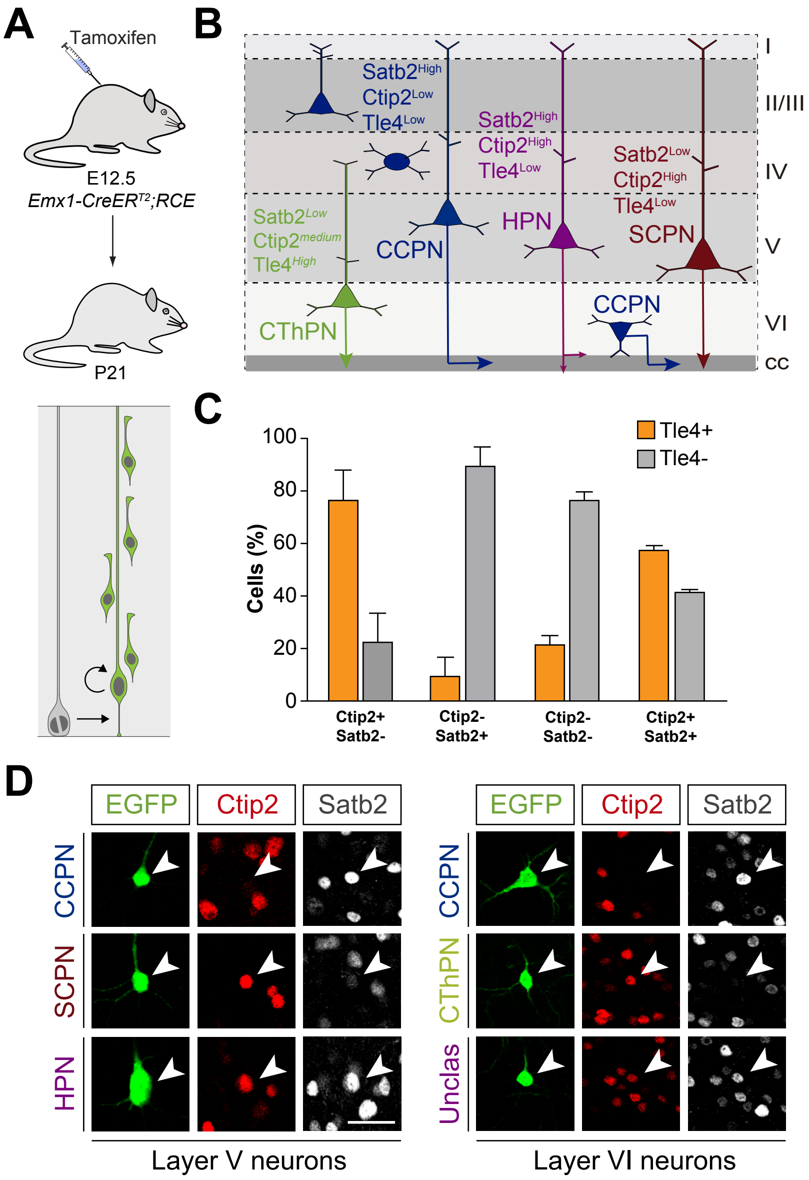

A
Tamoxifen
E12.5
Emx1-CreERT2;RCE
P21
B
Satb2High
Ctip2Low
Tle4Low
Satb2High
Ctip2High
Tle4Low
Satb2Low
Ctip2High
Tle4Low
Satb2Low
Ctip2medium
Tle4High
CCPN
HPN
SCPN
CThPN
CCPN
I
II/III
IV
V
VI
cc
C
Tle4+
Tle4-
Cells (%)
100
80
60
40
20
0
Ctip2+ Satb2-
Ctip2- Satb2+
Ctip2- Satb2-
Ctip2+ Satb2+
D
EGFP
Ctip2
Satb2
CCPN
SCPN
HPN
Layer V neurons
EGFP
Ctip2
Satb2
CCPN
CThPN
Unclas
Layer VI neurons

**Figure S5. Identification of pyramidal cell subclasses**

(A) Experimental paradigm. The bottom panel illustrates the expected labeling outcome of a neurogenic RGC division following inducible conditional reporter lineage tracing in *Emx1-CreER$^{T2}$;RCE* mice.
(B) Schematic of PC subclasses based on their laminar distribution and expression of specific markers.
(C) Fraction of Tle4+ and Tle4- layer VI PCs that express the transcription factors Satb2 and Ctip2. Data are represented as mean ± standard deviation.
(D) Pyramidal cell subclasses in lineages traced in *Emx1-CreER$^{T2}$;RCE* mice. Neurons are classified according to their laminar allocation and expression of Satb2 and Ctip2.
CCPN, cortico-cortical projection neuron; SCPN, subcortical projection neuron; HPN, heterogeneous projection neuron; CThPN, cortico-thalamic projection neuron.
Scale bar equals 35 µm.

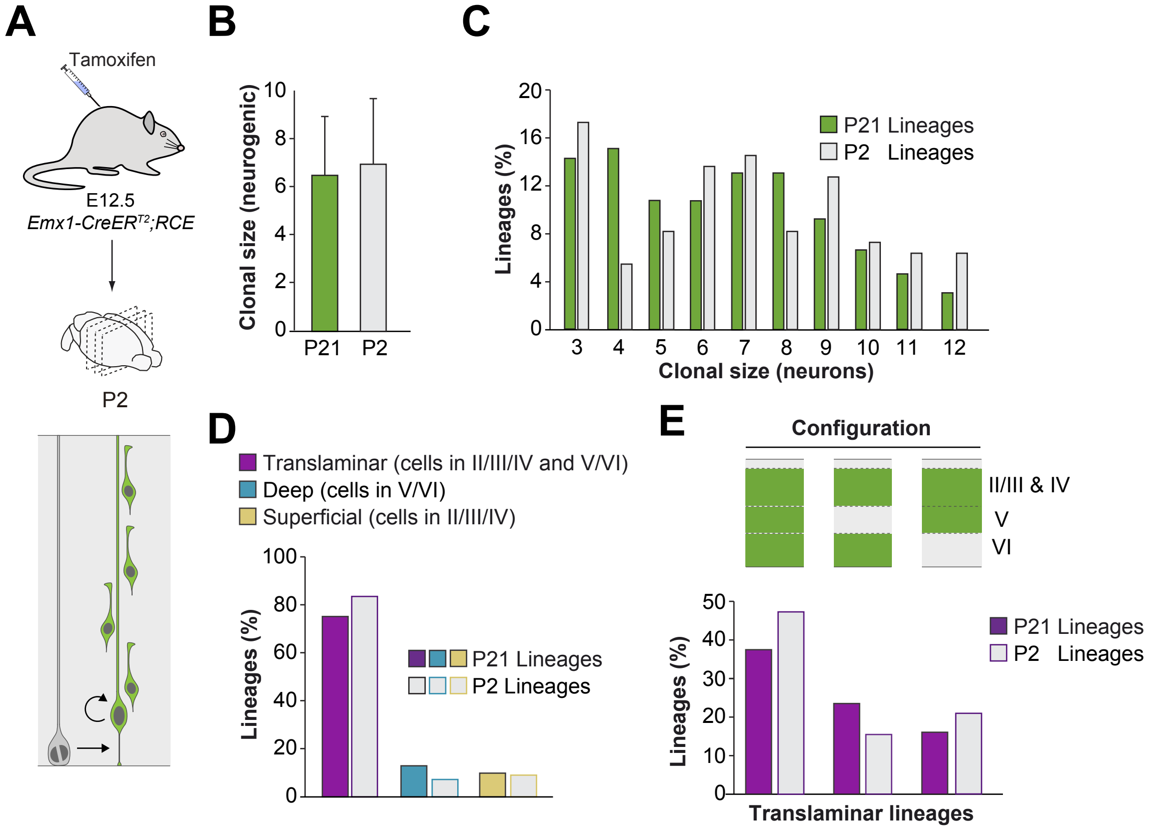
A
Tamoxifen
E12.5
Emx1-CreER^T2;RCE
P2
B
Clonal size (neurogenic)
10
8
6
4
2
0
P21
P2
C
Lineages (%)
20
16
12
8
4
0
P21 Lineages
P2 Lineages
Clonal size (neurons)
D
Translaminar (cells in II/III/IV and V/VI)
Deep (cells in V/VI)
Superficial (cells in II/III/IV)
Lineages (%)
100
80
60
40
20
0
P21 Lineages
P2 Lineages
E
Configuration
II/III & IV
V
VI
Lineages (%)
50
40
30
20
10
0
P21 Lineages
P2 Lineages
Translaminar lineages

**Figure S6. Subtle impact of pyramidal neuron cell death in final configurations of cortical neuron lineages**

(A) Experimental paradigm. The bottom panel illustrates the expected labeling outcome of a neurogenic RGC division following inducible conditional reporter lineage tracing in *Emx1-CreER$^{T2}$;RCE* mice.
(B) Clonal size of cortical lineages at P2 and P21 following low-dose tamoxifen in *Emx1-CreER$^{T2}$;RCE* mice at E12.5. Data are represented as mean ± standard deviation.
(C) Clonal size distribution of cortical lineages at P2 and P21 in inducible conditional reporter lineage tracing experiments. At this age, layers II/II and IV cannot be reliably distinguished and are consequently considered as a single cortical layer.
(D) Quantification of the fraction of translaminar, deep and superficial layer-restricted lineages at P2 and P21 in inducible conditional reporter lineage tracing experiments.
(E) Relative frequency (expressed as percentage over the total number of lineages) of the different laminar configurations (green and grey schemas) in inducible conditional reporter lineage tracing experiments at P2 and P21.

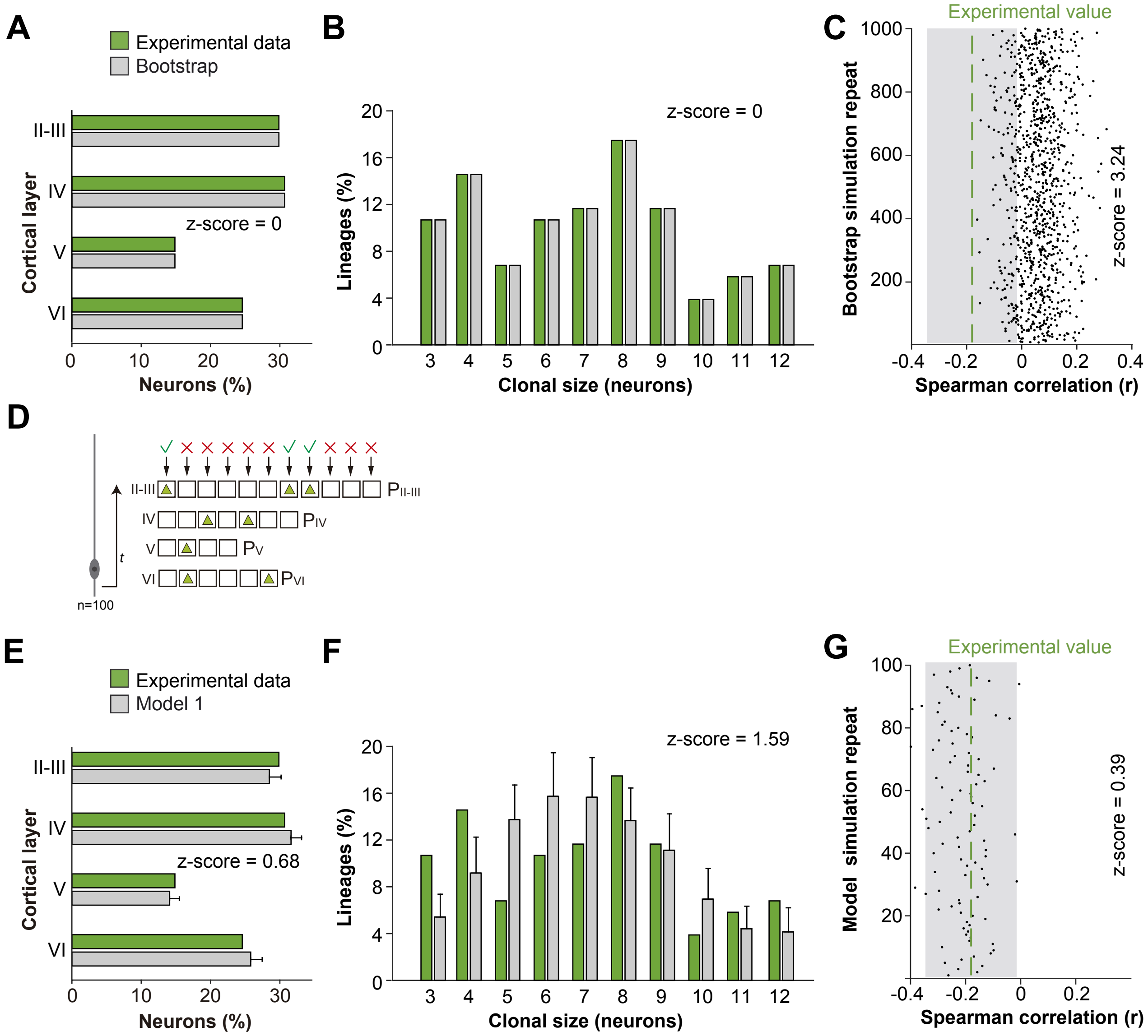
A
Experimental data
Bootstrap
II-III
IV
V
VI
Cortical layer
z-score = 0
Neurons (%)
B
Lineages (%)
z-score = 0
Clonal size (neurons)
C
Experimental value
Bootstrap simulation repeat
z-score = 3.24
Spearman correlation (r)
D
II-III
IV
V
VI
PII-III
PIV
PV
PVI
t
n=100
E
Experimental data
Model 1
Cortical layer
z-score = 0.68
Neurons (%)
F
Lineages (%)
z-score = 1.59
Clonal size (neurons)
G
Experimental value
Model simulation repeat
z-score = 0.39
Spearman correlation (r)

**Figure S7. A unique progenitor population cannot reproduce the observed diversity of cortical lineage configuration**

(A–C) Control parameters used to evaluate the approximation of random bootstrap modelling to the experimental dataset include the fraction of PCs in each cortical layer (A), the clonal size (B) and the spearman correlation (r) value for superficial and deep layer neurons (C) in bootstrapped lineages. Each dot in (C) represents an average r value for one simulation. The green line shows the experimental value; the shadow area around the experimental data represents a 95% confidence interval for the experimental value.
(D) Schematic representation of a mathematical model of cortical neurogenesis in which a single progenitor identity is modeled (Model 1).
(E–G) Control parameters used to evaluate the approximation of Model 1 to the experimental dataset include the fraction of PCs in each cortical layer (E), the clonal size (F) and the spearman correlation (r) value for superficial and deep layer neurons (G) in modeled lineages. Each dot in (G) represents an average r value for one simulation. The green line shows the experimental value; the shadow area around the experimental data represents a 95% confidence interval for the experimental value.
Histograms represent mean ± standard deviation. Z-scores represent the distance between experimental and simulated datasets for each parameter, which is calculated as the difference between the averages of model and experimental data divided by the standard deviation within model simulations (see Methods).

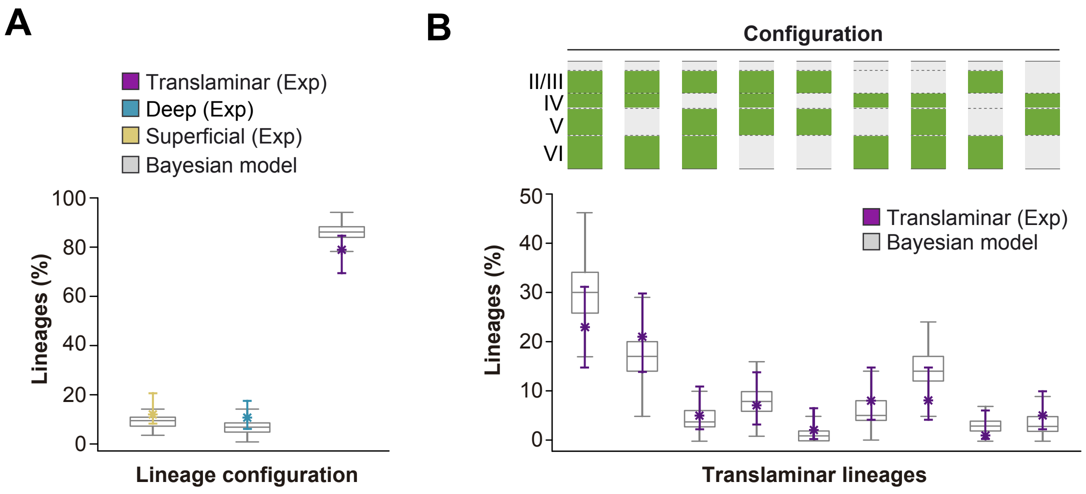
A
Translaminar (Exp)
Deep (Exp)
Superficial (Exp)
Bayesian model
Lineages (%)
100
80
60
40
20
0
Lineage configuration
B
Configuration
II/III
IV
V
VI
50
40
30
20
10
0
Lineages (%)
Translaminar (Exp)
Bayesian model
Translaminar lineages

**Figure S8. Bayesian modeling reproduces experimental lineage configurations**

(A) Distribution of translaminar, deep and superficial layer-restricted lineages predicted by the Bayesian model (expressed as percentage over all modelled lineages within a single simulation). Boxes represent variability obtained among 2000 posterior samples, colored stars and lines represent experimental values and 95% confidence intervals for experimental values, respectively.

(B) Relative frequency (expressed as percentage over all modeled translaminar lineages within a single simulation) of laminar configurations in experimental and Bayesian-modeled translaminar lineages. Grey boxes represent variability among 2000 posterior samples; colored stars and lines show experimental values and 95% confidence intervals for experimental values.

**Supplementary Table 1.** Summary of data and statistical analyses for Figures 1-6 and Supplementary Figures S1-S8

| **FIGURE 1** | Measurement | Values | N | Statistical | P value |
|---|---|---|---|---|---|
| Figure 1D | Number of neurons per lineage (mean ± SEM) | E9.5: 199.23 ± 28.83; E10.5: 59.62 ± 8.02; E11.5: 13.04 ± 0.94; E12.5: 6.14 ± 0.27; E14.5: 2.84 ± 0.17 | [Clones] E9.5, n = 13; E10.5, n = 21; E11.5, n = 51; E12.5, n = 73; E14.5, n = 32; [Brains] E9.5, n = 4; E10.5, n = 3; E11.5, n = 5; E12.5, n = 7; E14.5, n = 3 | | |
| Figure 1E | Fraction of one cell, two cells, and three or more cell lineages (percentage over total) | One cell: E9.5: 0%; E10.5: 0%; E11.5: 18.75%; E12.5: 39.15% | [Clones] E9.5, *n* = 13; E10.5, *n* = 21; E11.5, *n* = 64; E12.5, n = 166; [Brains] E9.5, n = 4; E10.5, n = 3; E11.5, n = 5; E12.5, n = 7; | | |
| | | Two cells: E9.5: 0%; E10.5: 0%; E11.5: 1.56%; E12.5: 16.46% | [Clones] E9.5, n = 13; E10.5, *n* = 21; E11.5, *n* = 64; E12.5, n = 166; [Brains] E9.5, n = 4; E10.5, n = 3; E11.5, n = 5; E12.5, n = 7 | | |
| | | Three or more cells: E9.5: 100%; E10.5: 100%; E11.5: 79.69%; E12.5: 44.39%; | [Clones] E9.5, n = 13; E10.5, *n* = 21; E11.5, *n* = 64; E12.5, n = 166; [Brains] E9.5, n = 4; E10.5, n = 3; E11.5, n = 5; E12.5, n = 7 | | |
| **FIGURE 2** | Measurement | Values | N | Statistical | P value |
| Figure 2E | Fraction of translaminar, deep, and superficial lineages (precentage over total) | Translaminar: 63.01%; Deep: 15.07%; Superficial: 21.92% | [Clones]: n = 73 [Brains]: n = 7 | | |
| | Number of neurons per lineage (mean ± Std) | Translaminar: 6.96 ± 2.38; Deep: 4.45 ± 0.93; Superficial: 4.94 ± 1.24% | [Clones]: n = 73 [Brains]: n = 7 | | |
| Figure 2I | Fraction of translaminar, deep, and superficial lineages (precentage over total) | Translaminar: 6.67%; Deep: 80%; Superficial: 13.33% | [Clones]: n = 30 [Brains]: n = 7 | | |
| Figure 2M | Fraction of lineages per cortical layer (precentage over total) | Layer II/III: 4.76%; Layer IV: 19.05%; Layer V: 17.46%; Layer VI: 58.73% | [Clones]: n = 63 [Brains]: n = 7 | | |
| **FIGURE 3** | Measurement | Values | N | Statistical | P value |
| Figure 3D | Fraction of translaminar, deep, and superficial lineages (precentage over total) | Translaminar: 91.05%; Deep: 6.6%; Superficial: 1.89% | [Clones]: n = 106 [Brains]: n = 28 | | |
| | Number of neurons per lineage (mean ± Std) | Translaminar: 8.05 ± 2.83; Deep: 4.00 ± 1.00 | [Clones]: n = 106 [Brains]: n = 28 | | |
| **FIGURE 4** | Measurement | Values | N | Statistical | P value |
| Figure 4E | Fraction of translaminar, deep, and superficial lineages (precentage over total) | Translaminar: 75.38%; Deep: 13.46%; Superficial: 11.15% | [Clones]: n = 260 [Brains]: n = 25 | | |
| | Number of neurons per lineage (mean ± Std) | Translaminar: 7.22 ± 2.37; Deep: 3.77 ± 1.29; Superficial: 4.90 ± 1.82 | [Clones]: n = 260 [Brains]: n = 25 | | |
| **FIGURE 5** | Measurement | Values | N | Statistical | P value |
| Figure 5E | Fraction of lineages containing 3 to 12 cells (percentage over total) | Translaminar: [3]: 3.46%; [4]: 8.08%; [5]: 8.85%; [6]: 9.23%; [7]: 11.54%; [8]: 11.54%; [9]: 8.85%; [10]: 6.15%; [11]: 4.62%; [12]: 3.08%; Deep: [3]: 8.08%; [4]: 3.08%; [5]:1.15%; [7]: 0.77%; [8]: 0.38%; Superficial: [3]: 2.69; [4]: 3.85%; [5]: 0.77%; [6]: 1.54%; [7]: 0.77%; [8]: 1.15%; [9]: 0.38% | [Clones]: n = 260 [Brains]: n = 25 | | |
| Figure 5F | Fraction of lineages in each configuration (percentage over total) | [II/III]: 1.92%; [IV]: 1.54%; [II/III-IV]: 7.69%; [V]: 1.54%; [VI]: 5.77%; [V-VI]: 6.15%; [II/III to VI]: 22.69%; [II/III-IV-VI]: 14.62%; [II/III-V-VI]: 10.38%; [II/III to V]: 7.31%; [II/III-V]: 5.38%; [IV-VI]: 4.62%; [IV to VI]: 3.85%; [II/III-VI]: 3.46%; [IV-V]: 3.08% | [Clones]: n = 260 [Brains]: n = 25 | | |
| Figure 5G | Number of lineages in each combination(total number) | [y-x] // [2-2]: 2; [2-3]: 1; [2-4]: 1; [3-3]: 3; [3-5]: 1; [3-6]: 1; [3-8]: 2; [4-2]: 3; [4-3]: 3; [4-4]: 3; [4-5]: 3; [4-6]: 1; [5-2]: 4; [5-3]: 3; [5-4]: 2; [5-5]: 3; [6-2]: 3; [6-3]: 2; [6-4]: 3; [6-5]: 4; [7-3]: 3; [7-4]: 2; [8-3]: 1; [8-4]: 1; [9-3]: 1; [10-2]: 1 | [Clones]: n = 57 [Brains]: n = 25 | | |
| Figure 5H | Fraction of lineages containing each sutype combination (precentage over total all-layer lineages) | [CCPN]: 23.08%; [CCPN; L5SCPN]: 7.69%; [CCPN; L6CThPN]: 11.54%; [CCPN; L5HPN]: 11.54%; [CCPN; L5HPN; L6CThPN]: 15.38%; [CCPN; L5SCPN; L5HPN]: 7.69%; [CCPN; L5SCPN; L6CThPN]: 15.38%; [CCPN; L5SCPN; L5HPN; L6CThPN]: 7.69% | [Clones]: n = 26 [Brains]: n = 25 | | |

| FIGURE 6 | Measurement | Values | N | Statistical | P value |
|---|---|---|---|---|---|
| Figure 6B | Fraction of cells per layer (percentage over total & mean percentage ± Std) | Experimental data, [II/III]: 29.86%; [IV]: 30.69%; [V]: 14.86%; [VI]: 24.58%; Model 2, [II/III]: 30.04 ± 1.82%; [IV]: 31.08 ± 1.85%; [V]: 14.19 ± 1.44%; [VI]: 24.70 ± 1.78% | Experimental, [Clones]: n = 103; [Brains]: n = 25; Model 2, [Clones]: n = 103 | | |
| Figure 6C | Fraction of lineages containing 3 to 12 cells (percentage over total & mean percentage ± Std) | Experimental data, [3]: 10.68%; [4]: 14.56%; [5]: 6.80%; [6]: 10.68%; [7]: 11.65%; [8]: 17.48%; [9]: 11.65%; [10]: 3.88%; [11]: 5.83%; [12]: 6.80%; Model 2, [3]: 11.46 ± 3.45%; [4]: 13.00 ± 3.49%; [5]: 12.14 ± 3.94%; [6]: 11.39 ± 2.96%; [7]: 11.47 ± 3.44%; [8]: 10.98 ± 3.24%; [9]: 8.85% ± 2.82%; [10]: 6.80 ± 2.95%; [11]: 5.05 ± 2.36%; [12]: 8.87 ± 3.27% | Experimental, [Clones]: n = 103; [Brains]: n = 25; Model 2, [Clones]: n = 103 | | |
| Figure 6E | Fraction of Translaminar, Deep, and Superficial lineages (precentage over total) | Experimental data (mean {I.C. 95%}): Translaminar: 77.67 {68.93-84.47}%; Deep: 10.68 {5.83-17.48}%; Superficial: 11.65 {6.8-19.42}%; Model 2 (median ± intercuartile distance), Translaminar: 78.57 ± 6.28%; Deep: 11.24 ± 3.91%; Superficial: 10.06 ± 5.14% | Experimental, [Clones]: n = 103; [Brains]: n = 25; Model 2, [Clones]: n = 103 | Fisher's exact test | p = 0.94 |
| Figure 6F | Fraction of lineages in each configuration (percentage over total) | Experimental data (mean {I.C. 95%}), [II/III to VI]: 22.33 {14.56-31.07}%; [II/III-IV-VI]: 20.39 {13.59-29.61}%; [II/III-V-VI]: 4.85 {1.94-10.68}%; [II/III to V]: 6.80 {2.91-13.59}%; [II/III-V]: 1.94 {0-6.09}%; [IV-VI]: 7.77 {3.88-14.56}%; [IV to VI]: 7.77 {3.88-14.56}%; [II/III-VI]: 0.97 {0-5.82}%; [IV-V]: 4.85 {1.94-19.71}%; Model 2 (median ± intercuartile distance), [II/III to VI]: 35.76 ± 8.06%; [II/III-IV-VI]: 20.24 ± 5.11%; [II/III-V-VI]: 4.20 ± 2.50%; [II/III to V]: 6.98 ± 4.45%; [II/III-V]: 1.15 ± 1.22%; [IV-VI]: 1.25 ± 1.45%; [IV to VI]: 3.53 ± 3.52%; [II/III-VI]: 2.41 ± 2.38%; [IV-V]: 0 ± 1.19% | Experimental, [Clones]: n = 103; [Brains]: n = 25; Model 2, [Clones]: n = 103 | Chi-square test | p = 0.18 |
| Figure 6G | Likelihood for each number of categories (percentage over total samples) | [1]: 0.2%; [2]: 0.75%; [3]: 82.50%; [4]: 12.45%; [5]: 0.5% | [Samples]: n = 200000 | | |
| Figure 6I | Fraction of cells per layer (percentage over total ± Std) | Fucci quantification, S1: [II/III]: 25.68 ± 2.13%; [IV]: 24.63 ± 0.63%; [V]: 16.61 ± 0.67%; [VI]: 33.09 ± 2.38%; Vi1: [II/III]: 30.39 ± 1.10%; [IV]: 24.31 ± 1.32%; [V]: 21.86 ± 1.22%; [VI]: 23.44 ± 1.58%; Model 2, S1: [II/III]: 24.43 ± 2.12%; [IV]: 24.46 ± 2.04%; [V]: 14.31 ± 1.27%; [VI]: 36.12 ± 2.24%; Vi1; [II/III]: 30.17 ± 2.03%; [IV]: 24.30 ± 1.74%; [V]: 22.13 ± 1.75%; [VI]: 23.40 ± 1.71% | [Brains]: n = 3 | | |
| Figure 6J | Probability of cell generation (mean) | Type 1: [PII/III]: 0.118; [PIV]: 0.078; [PV]: 0.041; [PVI]: 0.099; Type 2: [PII/III]: 0.045; [PIV]: 0.031; [PV]: 0.0199; [PVI]: 0.1499; Type 3: [PII/III]: 0.0180; [PIV]: 0.1638; [PV]: 0.0715; [PVI]: 0.0806 | [Samples]: n = 200000 | | |
| **Suppl. FIGURE 1** | Measurement | Values | N | Statistical | P value |
| Figure S1B | Fraction of lineages with one or two reporters (percentage over total) | One: 95.83%; Two: 4.17% | [Clones]: n = 144 [Brains]: n = 4 | | |
| **Suppl. FIGURE 2** | Measurement | Values | N | Statistical | P value |
| Figure S2D | Fraction of translaminar, deep, and superficial lineages (precentage over total) | Translaminar: 2.78%; Deep: 56.94%; Superficial: 40.28% | [Clones]: n = 72 [Brains]: n = 12 | | |
| **Suppl. FIGURE 3** | Measurement | Values | N | Statistical | P value |
| Figure S3C | Fraction of translaminar, deep, and superficial hemilineages (precentage over total) | Translaminar: 80.71%; Deep: 7.11%; Superficial: 12.18% | [Clones]: n = 106 [Brains]: n = 28 | | |
| **Suppl. FIGURE 4** | Measurement | Values | N | Statistical | P value |
| Figure S4D | Fraction of lineages containing 3 to 12 cells (percentage over total) | Translaminar, [4]: 5.48%; [5]: 19.18%; [6]: 9.59%; [7]: 4.11%; [8]: 6.85%; [9]: 9.59%; [10]: 2.74%; [11]: 2.74%; [12]: 1.37%; [14]: 1.37%; Deep, [3]: 2.74%; [4]: 4.12%; [5]: 6.85%; [6]: 1.37%; Superficial, [3]: 2.74; [4]: 5.48%; [5]: 6.85%; [6]:4.11%; [7]: 2.74 | [Clones]: n = 73 [Brains]: n = 7 | | |
| Figure S4E | Fraction of lineages in each configuration (percentage over total) | [IV]: 8.22%; [II/III-IV]: 13.7%; [V]: 1.37%; [VI]: 10.96%; [V-VI]: 2.74%; [II/III to VI]: 16.44%; [II/III-IV-VI]: 24.66%; [II/III to V]: 10.96%; [IV-VI]: 4.11%; [IV to VI]: 5.48%; [IV-V]: 1.37% | [Clones]: n = 73 [Brains]: n = 7 | | |
| Figure S4I | Fraction of lineages containing 3 to 12 cells (percentage over total) | Translaminar, [3]: 0.94%; [4]: 4.72%; [5]: 9.43%; [6]:12.26%; [7]: 14.15%; [8]: 17.92%; [9]: 12.26%; [10]: 3.77%; [11]: 5.66%; [12]: 5.66%; [13]: 0.94%; [14]: 0.94%; [15]: 1.89%; [21]: 0.94%; Deep, [3]: 1.89%; [4]: 3.77%; [6]: 0.94%; Superficial, [6]: 1.89% | [Clones]: n = 106 [Brains]: n = 28 | | |

| | | | | | |
|---|---|---|---|---|---|
| Figure S4J | Fraction of lineages in each configuration (percentage over total) | [II/III-IV]: 1.89%; [VI]: 6.6%; [II/III to VI]: 44.34%; [II/III-IV-VI]: 21.70%; [II/III-V-VI]: 1.89%; [II/III to V]: 9.43%; [IV-VI]: 3.77%; [IV to VI]: 9.43%; [IV-V]: 0.94% | [Clones]: n = 106 [Brains]: n = 28 | | |
| **Suppl. FIGURE 5** | Measurement | Values | N | Statistical | P value |
| Figure S5C | Fraction of cells expressing Tle4 (mean % ± Std) | C+/S-: [Tle4+]: 77.43 ± 12.46%; [Tle4-]: 22.57 ± 12.46; C-/S+: [Tle4+]: 10.19 ± 7.20%; [Tle4-]: 89.81 ± 7.20%; C-/S-: [Tle4+]: 22.39 ± 3.38%; [Tle4-]: 77.61 ± 3.38%; C+/S+: [Tle4+]: 58.21 ± 1.33%; [Tle4-]: 41.79 ± 1.33 | [Cells]: n = 1123 [Brains]: n = 2 | | |
| **Suppl. FIGURE 6** | | Values | N | Statistical | P value |
| Figure S6B | Number of neurons per lineage (mean ± Std) | [P21] : 6.51± 2.56; [P2]: 6.97 ± 2.76 | P21, [Clones]: n = 260; [Brains]: n = 25; P2, [Clones]: n = 110; [Brains]: n = 10 | Mann-Whitney test | p = 0.1501 |
| Figure S6C | Fraction of lineages containing 3 to 12 cells (percentage over total) | P21: [3]: 14.23%; [4]: 15.00%; [5]: 10.77%; [6]: 10.77%; [7]: 13.08%; [8]: 13.08%; [9]: 9.23%; [10]: 6.15%; [11]: 4.62%; [12]: 3.08%; P2: [3]: 17.27%; [4]: 5.45%; [5]: 8.18%; [6]: 13.63%; [7]: 14.55%; [8]: 8.18%; [9]: 12.72%; [10]: 7.27%; [11]: 6.36%; [12]: 6.36% | P21, [Clones]: n = 260; [Brains]: n = 25; P2, [Clones]: n = 110; [Brains]: n = 10 | Chi-square test | p = 0.1653 |
| Figure S6D | Fraction of translaminar, deep, and superficial lineages (precentage over total) | P21, Translaminar: 75.38%; Deep: 13.46%; Superficial: 11.15%; P2, Translaminar: 83.63%; Deep: 7.27%; Superficial: 9.09% | P21, [Clones]: n = 260; [Brains]: n = 25; P2, [Clones]: n = 110; [Brains]: n = 10 | Fisher's exact test | p = 0.1127 |
| Figure S6E | Fraction of lineages in each configuration (percentage over total) | P21: [II/III_&_IV to VI]: 36.78%; [II/III_&_IV-VI]: 22.99%; [II/III_&_IV to V]: 15.71%; P2: [II/III_&_IV to VI]: 47.27%; [II/III_&_IV-VI]: 15.45%; [II/III_&_IV to V]: 20.91% | P21, [Clones]: n = 260; [Brains]: n = 25; P2, [Clones]: n = 110; [Brains]: n = 10 | Chi-square test | p = 0.0994 |
| **Suppl. FIGURE 7** | Measurement | Values | N | Statistical | P value |
| Figure S7A | Fraction of cells per layer (percentage over total & mean percentage ± Std) | Experimental data, [II/III]: 29.86%; [IV]: 30.69%; [V]: 14.86%; [VI]: 24.58%; Bootstrap, [II/III]: 29.86 ± 0%; [IV]: 30.69 ± 0%; [V]: 14.86 ± 0%; [VI]: 24.58 ± 0% | Experimental, [Clones]: n = 103; [Brains]: n = 25; Bootstrap, [Clones]: n = 103 | | |
| Figure S7B | Fraction of lineages containing 3 to 12 cells (percentage over total & mean percentage ± Std) | Experimental data, [3]: 10.68%; [4]: 14.56%; [5]: 6.80%; [6]: 10.68%; [7]: 11.65%; [8]: 17.48%; [9]: 11.65%; [10]: 3.88%; [11]: 5.83%; [12]: 6.80%; Bootstrap, [3]: 10.68 ± 0%; [4]: 14.56 ± 0%; [5]: 6.80 ± 0%; [6]: 10.68 ± 0%; [7]: 11.65 ± 0%; [8]: 17.48 ± 0%; [9]: 11.65 ± 0%; [10]: 3.88 ± 0%; [11]: 5.83 ± 0%; [12]: 6.80 ± 0% | Experimental, [Clones]: n = 103; [Brains]: n = 25; Bootstrap, [Clones]: n = 103 | | |
| Figure S7E | Fraction of cells per layer (percentage over total & mean percentage ± Std) | Experimental data, [II/III]: 29.86%; [IV]: 30.69%; [V]: 14.86%; [VI]: 24.58%; Model 1, [II/III]: 28.49 ± 1.69%; [IV]: 31.61 ± 1.52%; [V]: 14.10 ± 1.40%; [VI]: 25.80 ± 1.65% | Experimental, [Clones]: n = 103; [Brains]: n = 25; Model 1, [Clones]: n = 103 | | |
| Figure S7F | Fraction of lineages containing 3 to 12 cells (percentage over total & mean percentage ± Std) | Experimental data, [3]: 10.68%; [4]: 14.56%; [5]: 6.80%; [6]: 10.68%; [7]: 11.65%; [8]: 17.48%; [9]: 11.65%; [10]: 3.88%; [11]: 5.83%; [12]: 6.80%; Model 1, [3]: 5.42 ± 1.95%; [4]: 9.18 ± 3.01%; [5]: 13.73 ± 2.98%; [6]: 15.73 ± 3.72%; [7]: 15.65 ± 3.39%; [8]: 13.65 ± 2.79%; [9]: 11.13 ± 3.11%; [10]: 6.95 ± 2.62%; [11]: 4.42 ± 1.93%; [12]: 4.15 ± 2.06% | Experimental, [Clones]: n = 103; [Brains]: n = 25; Model 1, [Clones]: n = 103 | | |
| **Suppl. FIGURE 8** | Measurement | Values | N | Statistical | P value |
| Figure S8A | Fraction of translaminar, deep, and superficial lineages (precentage over total) | Experimental data (mean {I.C. 95%}), Translaminar: 77.67 {68.93-84.47}%; Deep: 10.68 {5.83-17.48}%; Superficial: 11.65 {6.8-19.42}%; Bayesian model (median ± intercuartile distance), Translaminar: 85.43 ± 3.88%; Deep: 5.83 ± 3.88%; Superficial: 8.74 ± 2.91 | Experimental, [Clones]: n = 103; [Brains]: n = 25; Bayesian model, [Samples]: n = 200000 | | |
| Figure S8B | Fraction of lineages in each configuration (percentage over total) | Experimental data (mean {I.C. 95%}), [II/III to VI]: 22.33 {14.56-31.07}%; [II/III-IV-VI]: 20.39 {13.59-29.61}%; [II/III-V-VI]: 4.85 {1.94-10.68}%; [II/III to V]: 6.80 {2.91-13.59}%; [II/III-V]: 1.94 {0-6.09}%; [IV-VI]: 7.77 {3.88-14.56}%; [IV to VI]: 7.77 {3.88-14.56}%; [II/III-VI]: 0.97 {0-5.82}%; [IV-V]: 4.85 {1.94-19.71}%; Bayesian model (median ± intercuartile distance), [II/III to VI]: 29.13 ± 7.77%; [II/III-IV-VI]: 16.50 ± 5.83%; [II/III-V-VI]: 3.88 ± 2.91%; [II/III to V]: 7.77 ± 3.88%; [II/III-V]: 0.97 ± 1.94%; [IV-VI]: 4.85 ± 3.88%; [IV to VI]: 13.19 ± 4.95%; [II/III-VI]: 2.91 ± 1.94%; [IV-V]: 2.91 ± 2.91% | Experimental, [Clones]: n = 103; [Brains]: n = 25; Bayesian model, [Samples]: n = 200000 | | |

### METHODS

#### Experimental models

All transgenic mouse lines used in this study are listed on the key resources table. All adult mice were housed in groups and kept on a light/dark cycle (12/12 h) regardless of genotypes. Only time-mated pregnant female mice that have undergone in utero surgeries were house individually. Male and female mice were used in all experiments. In utero experiments where performed at different developmental stages that range from E9.5 to E14.5. For histological analyses, mice ages range from P2 to P30. All procedures were approved by King's College London or IST Austria and were performed under UK Home Office project licenses or in accordance with Austrian Federal Ministry of Science and Research license and European regulations (EU directive 86/609, EU decree 2001-486). The day of vaginal plug was considered as embryonic day (E) 0.5 and the day of birth as postnatal day (P) 0.

#### Retroviral infection for clonal labelling

Cre-dependent conditional retroviral stocks encoding EGFP and membrane-bound mCherry reporters (Ciceri et al., 2013) were produced as previously described (Tashiro et al. 2006). In brief, Moloney murine leukemia viruses were produced by transfecting HEK293T cells with retroviral plasmids (*rv::dio-eGfp* or *rv::dio-mCherry*, *pCMV-Vsvg*, and *pCMV-GAG-pol*) using lipofectamine 2000. Forty-eight hours post transfection the supernatant was collected, concentrated and purified by two sequential rounds of ultracentrifugation. The viral pellet was re-suspended in sterile PBS and stored in aliquots at -80°C. Green and red viral stocks were produced in the same plates and mixed before concentration by ultracentrifugation.

For *in utero* injections, pregnant females were deeply anesthetized with isoflurane and the abdominal cavity was incised to expose uterus. Conditional retroviruses were injected at low-titer into the telencephalic ventricles of E9.5, E10.5, E11.5, E12.5 and E14.5 mouse embryos using an ultrasound-guided imaging system (Visualsonic) coupled with a nanoliter injector, as previously described (Ciceri et al., 2013; Pla et al., 2006). Some experiments were performed using *rv::dio-eGfp* exclusively. After the procedure, the uterine horns were returned to the abdominal cavity and the wound was surgically sutured. The pregnant female was placed in a 32°C recovering chamber for 30 mins post-surgery before returning to standard housing conditions.

#### Inducible genetic clonal labeling

*Emx1-Cre$^{ERT2}$,RCE* and *Tbr2-Cre$^{ERT2}$*, *RCE* pregnant females received a single intraperitoneal injection of low dose (1 ng/kg) tamoxifen dissolved in corn oil at E12.5. MADM clones were generated as described previously (Beattie et al., 2017; Hippenmeyer et al., 2010). In brief, timed pregnant females were injected intraperitoneally with tamoxifen dissolved in corn oil at E12.5 at a dose of 2-3 mg/pregnant dam. Live embryos were recovered at E18.5-E19 through caesarean section, fostered, and raised for further analysis at P21.

#### Histology

Postnatal mice were perfused transcardially with 4% paraformaldehyde (PFA) in PBS and the dissected brains were fixed for 2 h at 4°C in the same solution. Brains were serially sectioned at 100 μm on a vibratome (VT1000S, Leica) or a freezing microtome (SM 2010R, Leica), and free-floating coronal sections were then subsequently processed for immunohistochemistry as previously described (Pla et al.,

2006). Primary and secondary antibodies used in the study are listed in key resource table.

#### Imaging

Images were acquired using fluorescence microscopes (DM5000B, CTR5000 and DMIRB from Leica or Apotome.2 from Zeiss) coupled to digital cameras (DC500 or DFC350FX, Leica; OrcaR2, Hamamatsu) with the appropriate emission filter sets or in inverted confocal microscopes (Leica TCS SP8 and Zeiss LSM800 Airyscan).

#### *In silico* modelling of cortical development

Modelling of progenitor behavior was performed using Matlab (Mathworks). To avoid overfitting variability that could correspond to differences in progenitor behavior across cortical areas, simulations were compared to lineages obtained from the primary somatosensory cortex using the *Emx1-CreER$^{T2}$;RCE* experimental dataset.

The similarity of model results to experimental data was assessed based on three control parameters: proportion of cells per layer, clonal size distribution and Spearman correlation (r) values for number of cells in upper versus lower layers. For each control parameter, we computed a normalized z-score measure by taking the difference between the experimental value and the average value across simulation repeats, and then dividing by the standard deviation across simulation repeats. The control parameter was considered to be consistent when the z-score value was less than 1. To generate randomly permuted cortical lineages, neurons observed in our *Emx1$^{CreER}$* ; RCE experimental dataset were permuted among lineages while maintaining their laminar identities. This operation was repeated 1000 times, providing average and standard deviation values that were then used to compare with the experimental dataset.

Probabilistic models 1 and 2 simulated 100 progenitors undergoing cell generation sequentially, following the *in vivo* inside-out pattern. In each layer*, in silico* progenitors took a number of stochastic decisions for neuron generation; at each decision, a new neuron could be generated, or alternatively, the chance could be skipped without neuron generation. Sequential generation of neurons thus used the following parameters. Progenitor capacity was defined as the maximum number of neurons that can be generated by a single progenitor, and had a fixed value of 12 cells, matching the experimentally cap. Number of opportunities per layer was established as the maximum number of cells found in a particular layer across all experimental lineages observed with all techniques. This parameter corresponds to the number of stochastic decisions available to the progenitor and reflects the size of the temporal "window" within which a progenitor can generate neurons in a given layer. Probability of cell generation, also layer-specific, gave the likelihood that a neuron is actually generated at each decision point. Simulations were repeated 100 times. Lineages with less than three cells were discarded for analysis.

For each model, the set of laminar division probabilities was adjusted to find the closest possible fit of lineage configurations to experimental data, according to the control parameters. Model 1 used a unique progenitor, i.e., a single set of laminar division probabilities. Model 2 incorporated two progenitor populations and was fit by varying both the relative size of the two populations and the values of their division probabilities, including how probabilities varied across layers.

#### Bayesian inference of progenitor types

In order to perform statistical inference on the number of progenitor types required to explain the distribution of lineages throughout the cortical layers, we employed a statistical model where $N$ observed lineages are grouped in $K$ progenitor types. Each type $t_{1:K}$ is associated to a vector of four probabilities $p_t = \{p_t^{(II/III)}, p_t^{(IV)}, p_t^{(V)}, p_t^{(VI)}\}$

representing probabilities for laminar occupancy. We assume that each observed lineage can be assigned to a unique progenitor type based on its occupancy distribution. Progenitor types are associated with frequencies $f_t$ , reflecting how likely is a lineage to belong to type *t.* The parameters $p$'s and $f$'s for each type as well as the number of types $K$ required can be obtained using Bayesian inference according to the Bayes' theorem

$$P(t_{1:N}, p_{1:K}, f_{1:K}|S) = \frac{\overbrace{P(t_{1:N}, S|p_{1:K}, f_{1:K})}^{likelihood} \cdot \overbrace{P(p_{1:K}; f_{1:K})}^{prior}}{\underbrace{P(S)}_{marginal\ likelihood}}$$

where $S$ is the occupancy matrix with the elements $S_{ij}$ representing the number of neurons $i$ found in layer $j$. The Bayes' theorem provides the posterior distribution of the model parameters $p$'s and $f$'s as well as the type of each progenitor conditional to the observations.

Our statistical model can be viewed as the following two-step generative process:

1. Each lineage $i$ is assigned to a progenitor type $t_i$ drawn independently from a categorical distribution with frequencies $f$'s.

*2.* The occupancy vector $S_{ij}$ of each lineage $i$ at each layer $j$ is drawn from a binomial distribution $Binomial\left(p_{t_i}^{(j)}; N_{max}\right)$where $N_{max}=20$ is the maximum number of cells (per progenitor) that can occupy each layer.

The likelihood of a given configuration $\{t_{1:K}, S\}$ reads

$$P(t_{1:K}, S|p_{1:K}, f_{1:K}) = \prod f_t^{n_t} \prod p_{t_i}^{S_{ij}} \left(1 - p_{t_i}^{N_{max}-S_{ij}}\right)$$

To perform Bayesian inference, we have chosen a Dirichlet prior distribution on the type frequencies $f_{1:K}$ and Beta distributions as priors on the occupancy probabilities $p_{1:K}$. To draw samples of model parameters and progenitor types from the posterior distribution we have implemented a Gibbs sampler (custom code written in C++ available upon request) which combines data likelihood and prior distributions to explore the parameter space efficiently. Within the Montecarlo sampler we have also allowed bidirectional transitions (increase or decrease) in the number of progenitor types controlled by a Metropolis-Hastings acceptance rule based on likelihood ratios between different number of types.

#### Quantification of cell distribution and clonal spatial configuration

In all the experiments, brain sections were sequentially analyzed in rostral-to-caudal order and pyramidal cell (PC) clones throughout the entire neocortex were identified as sparse, spatially separated cell clusters. The boundaries between cortical layers were traced based on nuclear (DAPI) staining and the laminar position of each cell was recorded accordingly. Pyramidal cell clones were classified as translaminar, deep layer- and superficial layer-restricted lineages according to the laminar position of the constituent neurons. Cortical areas were identified based on the reference atlas of adult mouse brain (Allen Brain Atlas; http://www.brain-map.org). In *Emx1-Cre$^{ERT2}$;MADM* experiments, lineages derived from symmetric divisions (defined as lineages with three or more cells expressing both reporters) were excluded. In *Emx1-Cre$^{ERT2}$;RCE* experiments, lineages derived from symmetric divisions (defined as lineages containing more than twelve neurons) were excluded. Lineages containing one or two cells were also excluded from *Emx1-Cre$^{ERT2}$;MADM* and *Emx1-Cre$^{ERT2}$;RCE* experiments.

#### Classification of pyramidal cell subclasses

Brain sections were stained with PC markers and classified based on the relative expression of the transcription factors Ctip2 and Satb2 in four main subclasses: Corticocortical projection neurons (CCPN), subcerebral projection neurons (SCPN), corticothalamic projection neurons (CThPN) and heterogeneous projection neurons (HPN). This last type was defined as layer V cells expressing both Ctip2 and Satb2, as described recently (Harb et al., 2016). Images were captured using a confocal microscope and analyzed with a custom algorithm written in Matlab (Mathworks). In brief, cell nuclei were segmented using the disk morphological function based on size and thresholds of fluorescence intensity over background. Cells were categorized as expressing high or low levels of the transcription factors Ctip2 and Satb2 and further subclassified as CCPN ($Ctip2^{Low}/Satb2^{High}$), SCPN ($Ctip2^{High}/Satb2^{Low}$) or HPN ($Ctip2^{High}/Satb2^{High}$). To distinguish between CThPN from CCPN in layer VI we used the following criteria: CCPN ($Ctip2^{Low}/Satb2^{High}$ or $Ctip2^{Low}/Satb2^{Low}$) or CThPN ($Ctip2^{High}/Satb2^{Low}$). This allowed the classification of layer V and layer VI cells based on the same set of markers. We verified these criteria by staining brain sections for the transcription factor Tle4 (Figure S6), a specific marker of cortical CThPN identity (Molyneaux et al., 2015). Layer VI cells expressing high levels of both transcription factors where not classified, and lineages containing those cells were excluded from the quantification.

#### Quantification of relative laminar ratios of pyramidal cells

To quantify PC densities in different cortical layers, $Nex^{Cre/+}$ mice were crossed with *Fucci2aR* reporter mice (Mort et al., 2014). The density of red nuclei in each cortical layer was quantified from five representative serial sections of the primary somatosensory and visual cortices. Z-stacks where then 3D-reconstructed and quantified using Imaris 8.1.2 (Bitplane).

#### Statistical tests

A summary of data and statistical analyses can be found in Table S1. Error bars in all graphs indicate standard deviation (std) unless otherwise stated. Comparisons of distributions over fractions of a total (as in Figure 6E and F and Figure S5C–E) were analyzed using Fisher's exact or Chi-square tests. Average clonal sizes between lineages were analyzed using U-Mann Whitney test. All statistical tests are specified in the figure legends.

#### Data and software availability

Custom-written MatLab (Mathworks, USA) codes used for quantification are available on request.
